# Bacterial Receptors but Not Anti-Phage Defense Mechanisms Determine Host Range for a Pair of *Pseudomonas aeruginosa* Lytic Phages

**DOI:** 10.1101/2024.04.30.591980

**Authors:** Dea M. Müller, Julie D. Pourtois, Minyoung K. Kim, Bobby Targ, Elizabeth B. Burgener, Carlos Milla, Maryam Hajfathalian, Hemaa Selvakumar, Joel Berry, Max Hopkins, Robert McBride, Jyot D. Antani, Paul E. Turner, Jonathan L. Koff, Paul L. Bollyky

## Abstract

Limited phage host range remains one of the obstacles to the widespread use of phage therapy against bacterial infections. Here, we perform a genome-wide association study (GWAS) using *Pseudomonas aeruginosa* clinical isolates collected from people with cystic fibrosis (pwCF) to identify bacterial genes associated with resistance or susceptibility to two lytic phages, OMKO1 and LPS-5, recently used in a clinical trial in pwCF. Results were validated with transposon mutagenesis experiments and functional assays. Genes associated with flagellum assembly and lipopolysaccharide biosynthesis are essential for infection by OMKO1 and LPS-5, respectively, consistent with functional studies implicating these molecules as receptors for these phages. Notably, the presence of bacterial genes encoding phage defense mechanisms is not predictive of phage susceptibility. Instead, the relative abundance of defense elements is associated with the number of temperate phages within bacterial genomes. Together, our findings highlight the central role of receptors in determining phage host range.

## Introduction

The emergence of antimicrobial-resistant (AMR) pathogens is a global health concern, requiring the development of innovative therapeutic strategies^1^. Among these resistant pathogens, the gram-negative bacterium *Pseudomonas aeruginosa (Pa)* is particularly problematic in the context of chronic lung infections in people with cystic fibrosis (pwCF)^2^ as well as in other settings^3,4^.

There is considerable interest in developing bacteriophages (phages) as therapeutic agents in the fight against AMR bacteria, including in pwCF^5^. Lytic bacteriophages are viral parasites of bacteria that use the bacterial cell to replicate. They adsorb and infect host bacteria via cell receptors, replicate using cellular machinery, and cause host cell lysis to release new phage particles enabling the initiation of new infection cycles in nearby cells^6,7^. Despite promising results^8–13^, therapeutic success is inconsistent^14–18^. While the reasons for these failures are multifactorial, the limited host range of most phages remains a fundamental challenge to scalable phage treatment of bacterial pathogens.

Phage receptor specificity is known to influence host range^19–21^. For a successful phage infection, the phage must first adsorb (bind) onto one or more specific receptors on the cell surface, such as proteins, protein complexes or sugar moieties on LPS, or other membrane structures. This results in many phages exhibiting a narrow host range, usually a subset of strains (genotypes) within the target bacterial species^22^. While this specificity can contribute to the safety of phage therapy because of limited infection of non-target cells, it also creates challenges for the development and application of phage therapy at a larger scale. Substantial effort is dedicated to engineering phages and increasing their host range, and to simplifying the pipeline from diagnosis to treatment^23^.

Bacterial resistance to phages is also implicated in phage host range. Bacteria and phages continuously evolve in a co-evolutionary arms race: bacteria acquire phage resistance mechanisms^24^, while phages may evolve counterstrategies in response^25^. Bacterial resistance mechanisms, including innate mechanisms (e.g., Restriction-Modification systems (RM) and toxin-antitoxins systems^26–28^) and adaptive mechanisms (e.g., CRISPR elements and CRISPR-associated (cas) genes^29,30^), protect against phage predation. It has been shown that 30% of known *Pa* strains possess a type I-F CRISPR-Cas system, 6% a type I-E system and only very few also carry the type I-C CRISPR type^26^. However, the extent to which these defense systems contribute to therapeutic failure is unclear, as many of the identified mechanisms (e.g., abortive infection (Abi) systems, RM systems, and CRISPR) are highly specific, often targeting only a limited range of phages. Addressing the biological limitations of phage infection remains a fundamental research focus and an essential step towards therapeutic success.

In this work, we investigated phage susceptibility patterns in two distinct collections of 80 clinical *Pa* isolates from pwCF, tested separately against OMKO1^31^ and LPS-5, two phages recently included in a human clinical trial of phage therapy (NCT04684641). We first performed a Genome-Wide Association Study (GWAS) analysis to identify bacterial genes associated with phage resistance or susceptibility. We then sought to validate the results using a transposon mutagenesis library screening and functional analyses. Finally, we evaluated the association between the presence of phage resistance mechanisms in the bacterial genome and phage susceptibility and prophage presence.

## Results

### Flagella-associated genes are significantly associated with OMKO1 phage susceptibility

A lytic plaque assay was conducted to determine OMKO1 (Figure 1A) and LPS-5 susceptibility in two distinct sets of 80 clinical *Pa* isolates (Figure 1B, Supplementary Table 1). These phages exhibit strong therapeutic potential, possess a relatively broad host range, and have been studied in the context of antibiotic synergy^31^ and inhaled phage therapy to mitigate virulence^16^.

**Figure 1:**
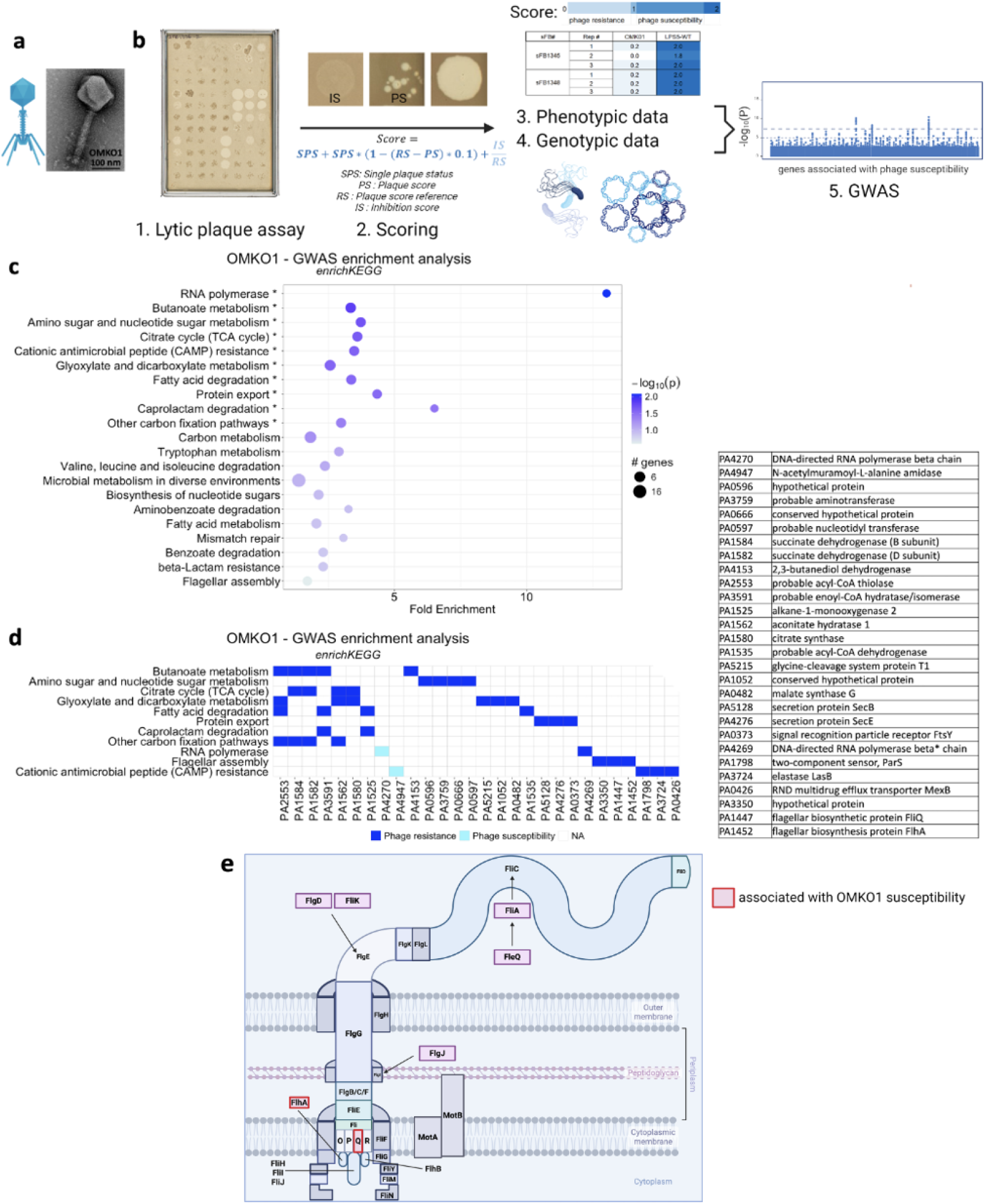
A GWAS study reveals genes and pathways associated with susceptibility to OMKO1. **A**, Transmission electron microscopy (TEM) image of the first phage OMKO1. **B**, Plaque assessment during the lytic plaque assay included single plaque status, plaque score of the sample and reference, and inhibition score. These factors collectively contributed to the derived score, which was used alongside genomic data for the GWAS. **C**, Functional enrichment analysis. Each bubble represents a specific functional category, with the size of the bubble indicating the number of associated genes, and the color intensity reflecting the statistical significance level of the enrichment. Significant pathways are labelled with a star. **D**, Genes associated with high phage susceptibility (light blue) or resistance (dark blue). **E**, Schematic representation of flagella and essential structural proteins. Proteins identified through the above described experiments as associated with OMKO1 phage susceptibility are highlighted in red.

To quantify plaquing, we developed a scoring system involving several parameters including single plaque status (SPS), plaque score of the sample (PS) and reference (RS), and inhibition score (IS), as described in detail in the methods section. These parameters collectively contributed to the derived score for each clinical isolate, ranging from 0 to 2. Scores between 0 and 1 indicate phage resistance or inhibition of bacterial growth, while scores between 1 and 2 indicate phage susceptibility.

This phenotypic data, combined with the whole-genome sequences of the clinical isolates, was used to perform a De Bruijn Genome-Wide Association Study (DBGWAS) to identify genes associated with either phage susceptibility or resistance. Quantile–quantile (QQ) plots were generated for diagnostic analysis (Supplementary Figure 1).

We then performed a functional enrichment analysis to identify pathways associated with phage infection. We found that genes for flagellar assembly as well as metabolic pathways were enriched in the set of genes associated with phage susceptibility, although not all pathways showed significant enrichment (Figure 1C). We then differentiated between genes from these pathways associated with phage susceptibility and those associated with phage resistance. Notably, genes *fliQ, flhA* and the hypothetical protein PA3350 in the flagellar assembly group are highlighted as important for OMKO1 phage resistance (Figure 1D). The flagellum variants were mostly located on the intracellular portion of the flagellum (Figure 1E).

To further validate the role for flagellum in OMKO1 infection, we conducted a transposon mutagenesis (Tn) assay (Figure 2A). Out of 49 identified genes that were significantly associated with OMKO1 phage susceptibility at a Multiplicity of Infection (MOI) of 0.1 (Fisher’s test, p-value < 0.05), 28 genes were involved in flagellum structure and function (Figure 2B, Supplementary Figure 2). We found that significant functional categories in the functional enrichment analysis are linked to flagella assembly, flagellum-dependent cell motility and flagellum organization (PANTHER, p-value < 0.05) (Figure 2C). OMKO1 infection was associated with many flagella genes encoding proteins across the physical flagellum structure, from cytoplasmic to extracellular components (Figure 2D). Similar results were obtained at MOI 20 (Supplementary Figure 3).

**Figure 2:**
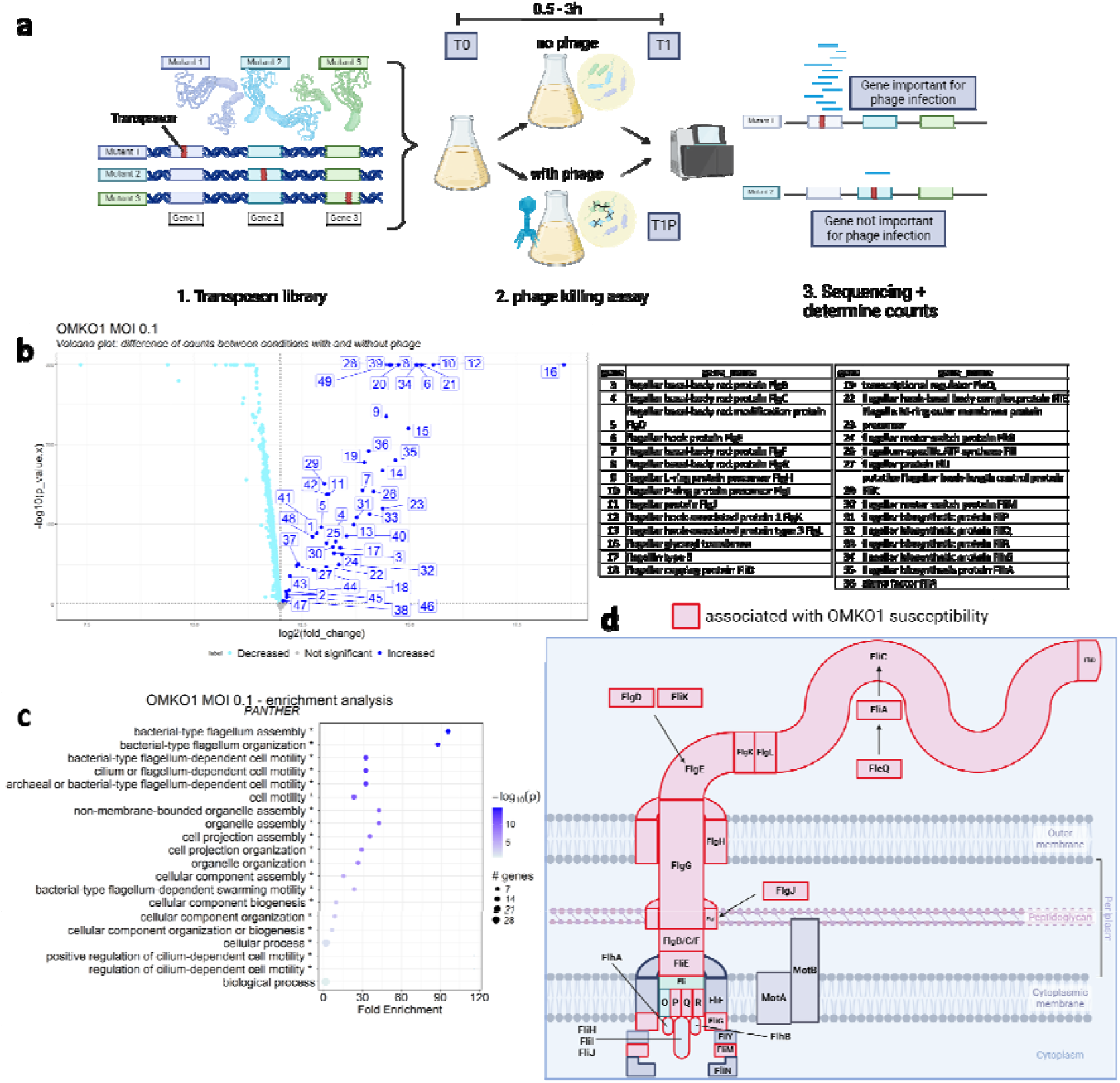
Transposon Mutagenesis Experiments show that Flagella-Associated Genes are required for OMKO1 Phage Infection. **A**, In the transposon mutagenesis library, a distinct gene in each mutant strain was disrupted via a transposon insertion. Following co-cultivation with phages, transposons were sequenced and quantified. Abundant transposon counts indicated bacterial survival, suggesting that the disrupted gene played a protective role against phage infection. This allowed the identification of genes implicated during phage infection. Conversely, minimal or absent transposon counts signified bacterial death and that the disrupted gene was not essential for phage infection. **B**, p-values for difference in gene counts (fold change) between conditions with and without OMKO1 phage at MOI 20. Each point represents a gene, with the x-axis showing the log2 fold change and the y-axis showing p-values for Fisher’s test. Significant increase in gene count is showed in dark blue, significant decrease in bright blue and non significant genes in gray. Genes showing a significant increase in counts in the presence of phage, indicative of their importance in phage infection, are labelled. (The complete list of significant genes can be found in Supplementary Figure 2) **C**, Functional enrichment analysis, where each bubble represents a functional category. The size of each bubble corresponds to the number of associated genes, and the colour intensity indicates the significance level. Significant pathways are labelled with a star. Functional enrichment analysis using PANTHER enables a detailed examination. Significant functional categories are associated with flagellum-dependent cell motility and flagellum assembly, particularly for the phage OMKO1. **D**, Schematic representation of flagella and essential structural proteins. Proteins identified through Tn mutagenesis experiments as associated with OMKO1 phage susceptibility are highlighted in red.

We further validated the involvement of the functional flagellum as a receptor for phage OMKO1 by conducting a suppression assay assessing phage infection of OMKO1 for 7 clinical isolates (Figure 3A) and evaluating the functionality of the flagellum in these clinical isolates with a swimming assay (Figure 3B). We found a significant relationship between suppression ratio and swimming area (t-test, p = 0.004, R^2^ = 0.81).

**Figure 3:**
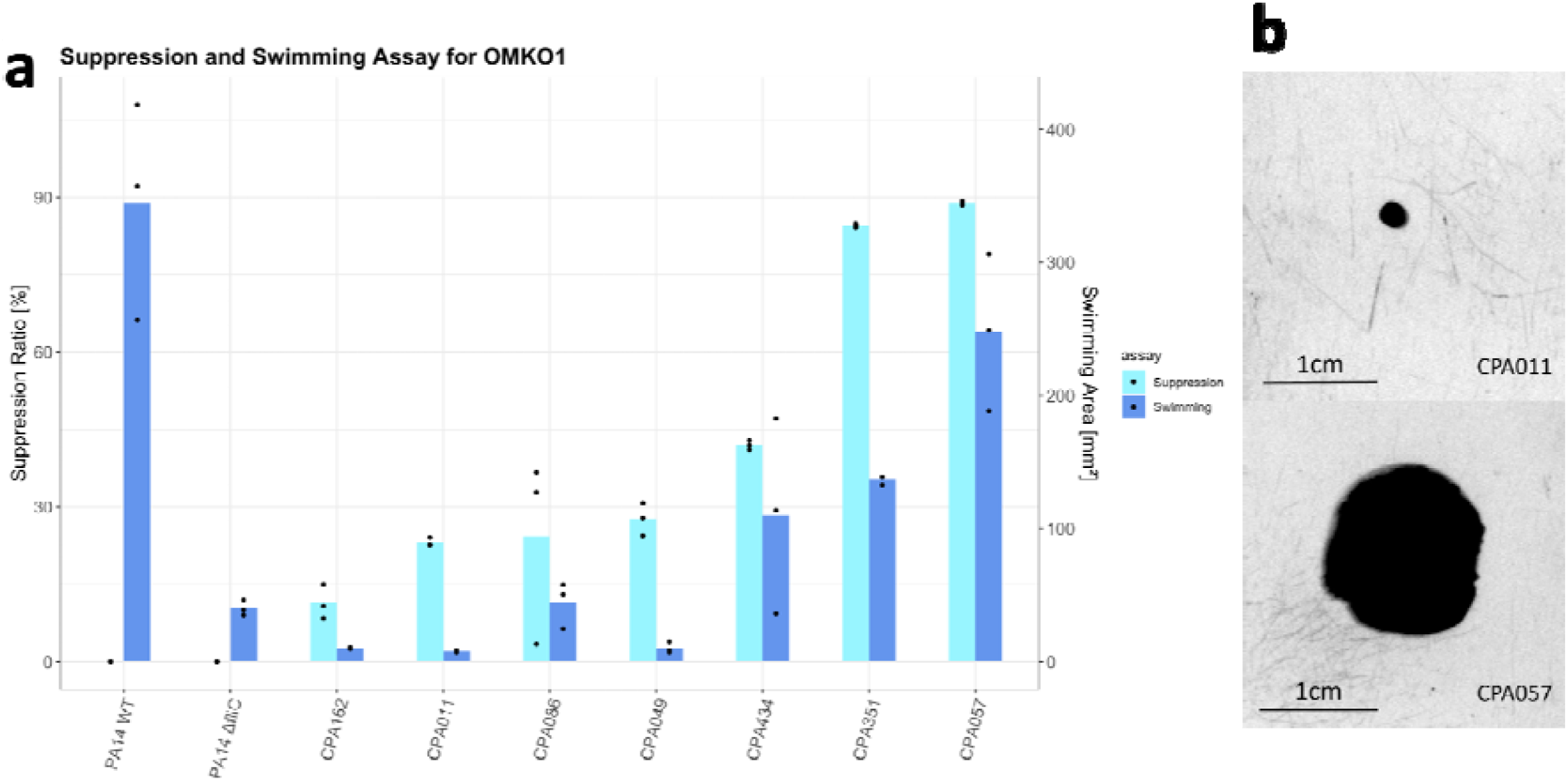
The presence of a functional bacterial flagellum is associated with susceptibility to OMKO1 infection. **A**, Phage suppression (Suppression ratio) was assessed for clinical Pseudomonas isolates (CPA). The suppression ratio was calculated as the area under the curve (AUC) for the non-phage-treated condition minus the phage-treated condition, divided by the AUC of the non-treated sample. Swimming assays for various CPAs. Growth areas similar to those of the Δ*fliC* strain indicate non-functional flagella, while swimming behavior comparable to that of the WT strain indicates functional flagella. We found a significant relationship between suppression ratio and swimming area (t-test, p = 0.004, R^2^ = 0.81). Phage suppression for WT and Δ*fliC* PA14 was not tested in this assay. **B**, Large colony growth area indicates the presence a functional flagellum allowing swimming (CPA057), whereas small colony growth area indicates a non-functional flagellum (CPA011).

Consistent with this role for flagella as a receptor for OMK01, we identified mutations in flagella genes from isolates resistant to OMKO1. The largest differences were observed in the genes *fliC, fliD, flgK*, and *flgL*. Sequence similarity for these genes was clustered according to percent identity within resistant and susceptible strains, and consensus sequences for both resistant and susceptible strains were generated (Supplementary Table 2). The pairwise identity of the consensus sequences was assessed (*fliC*=52.7%, *fliD*=only resistant strains had a blast output, *flgK*=83.4%, *flgL*=72.6%). Additionally, single mutations were identified in other flagella genes, but no associations to phage infection could be identified with the current sample size (Supplementary Table 3).

### LPS genes are associated with LPS-5 phage infection

We conducted a functional enrichment analysis from the GWAS data to identify enriched pathways in genes associated with LPS-5 phage (Figure 4A) susceptibility. We found several metabolic pathways that were enriched and relevant to phage susceptibility, including those involved in lipid metabolism (Figure 4B). We then differentiated between genes from these pathways associated with high phage susceptibility and those associated with phage resistance (Figure 4C).

**Figure 4:**
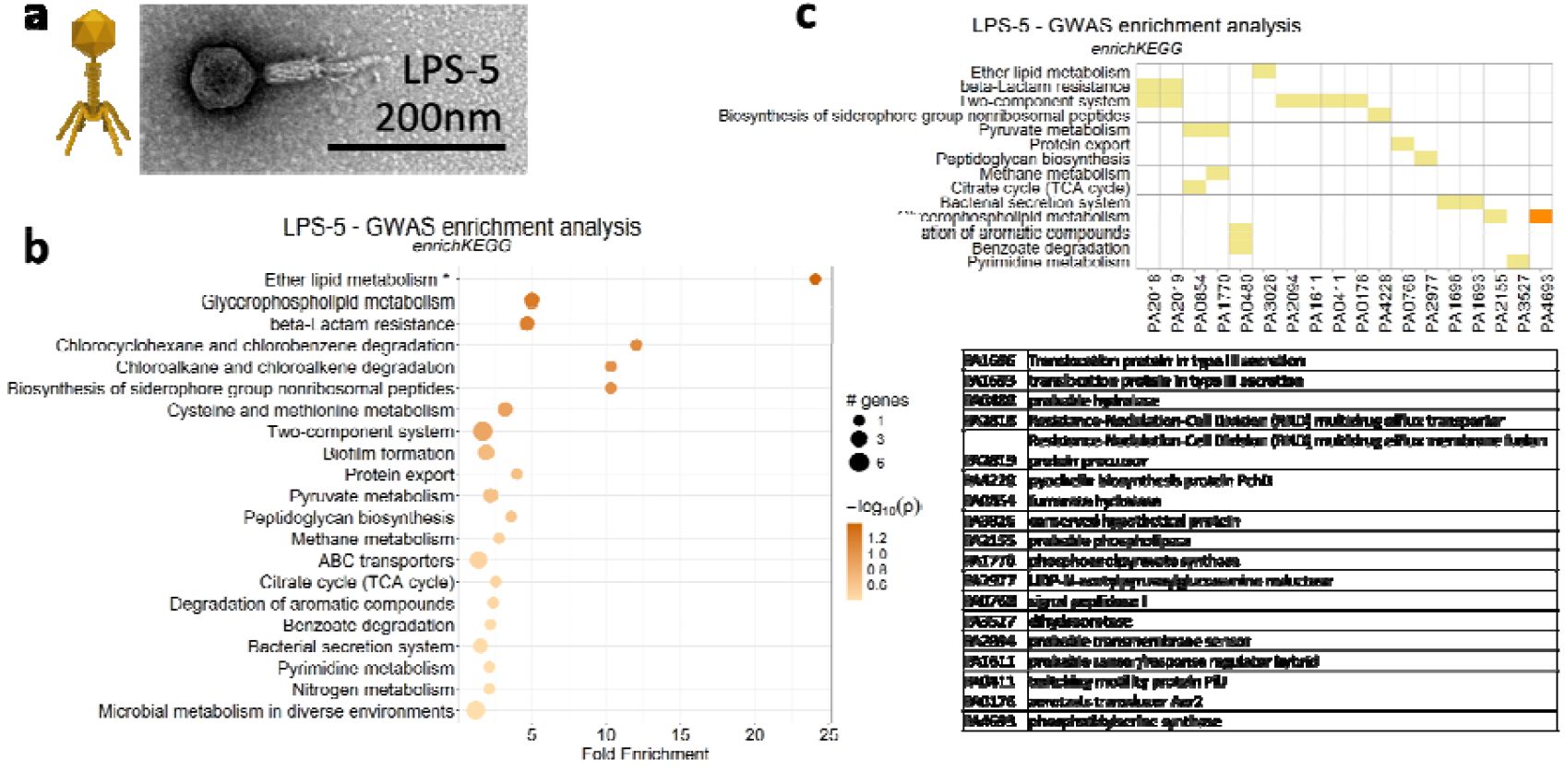
Lipid metabolism pathway is associated with LPS-5 susceptibility. **A**, TEM image of LPS-5 phage. **B**, Functional enrichment analysis. Each bubble represents a specific functional category, with the size of the bubble indicating the number of associated genes, and the color intensity reflecting the statistical significance level of the enrichment. Significant pathways are labelled with a star. **C**, Genes associated with high phage susceptibility (yellow) or resistance (orange).

To better identify genes involved in LPS-5 infection, we then performed a Tn mutagenesis assay. Using a MOI of 0.1, 29 genes were found to be significantly associated with LPS-5 infection (Fisher’s test, p-value < 0.05), including four genes related to lipopolysaccharide (LPS) biosynthesis (Figure 5A, Supplementary Figure 4). Enrichment analysis using PANTHER revealed only cellular metabolic process as a significantly enriched pathway but identified many pathways related to LPS function showing a trend of enrichment for the phage LPS-5 (Figure 5B). Some encoded proteins were involved in LPS biosynthesis pathways (Figure 5C). Similar results were obtained at MOI 20 (Supplementary Figure 5).

**Figure 5:**
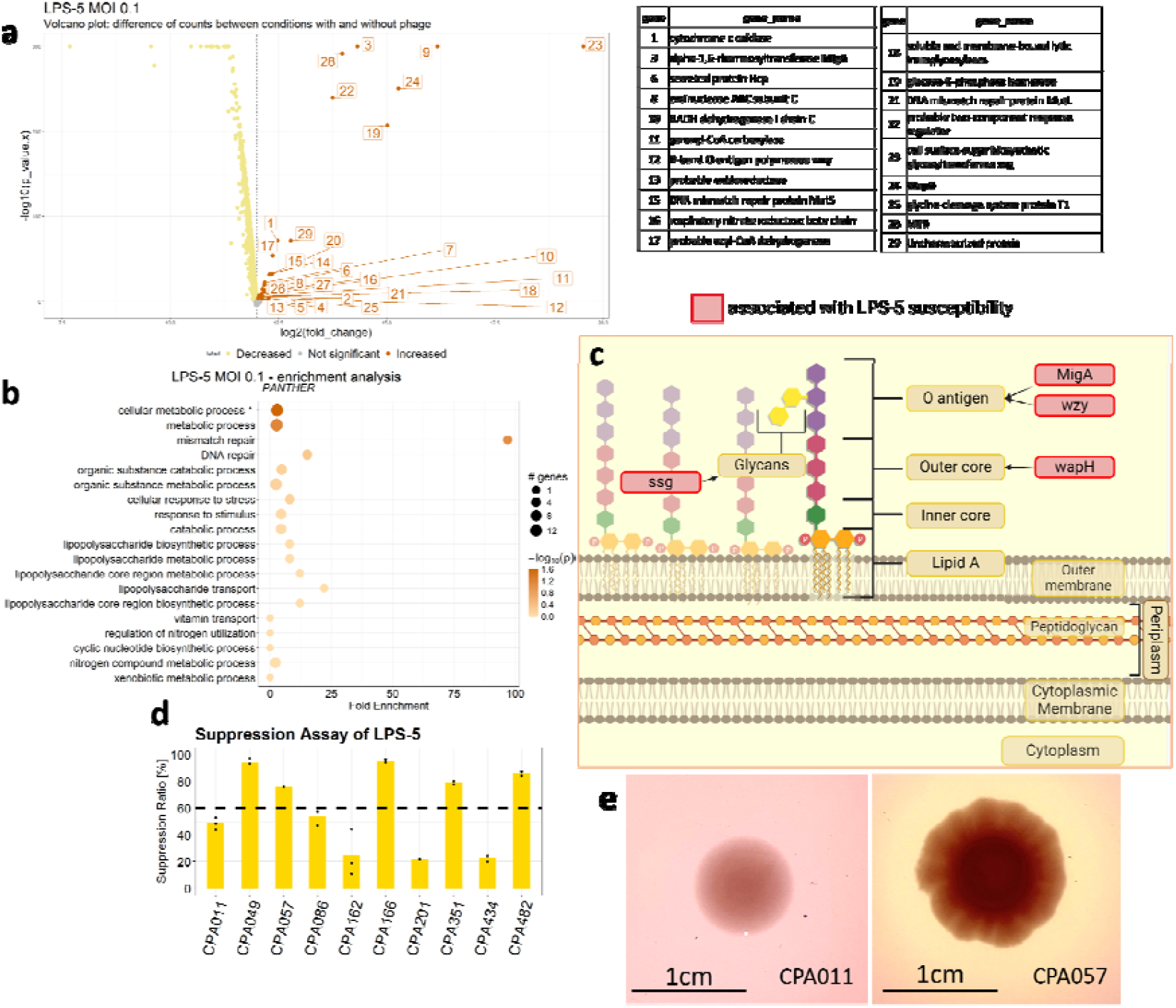
Identification of Genes associated with LPS-5 Phage Infection through Transposon Mutagenesis Experiments. **A**, This volcano plot highlights genes with a significant difference in counts (fold change) between conditions with and without LPS-5 phage at MOI 20. Each point represents a gene, with the x-axis showing the log2 fold change and the y-axis representing statistical significance. Significant increase in gene count is showed in orange, significant decrease in yellow and non significant genes in gray. Notably, several genes showing a significant increase in counts in the presence of phage, indicative of their importance in phage infection, are associated with LPS. (The complete list of significant genes can be found in Supplementary Figure 4.) **B**, Functional enrichment analysis using PANTHER. Several functional categories are associated with LPS and LPS biosynthesis, particularly for the phage LPS. **C**, Schematic representation of LPS and related pathways. Proteins identified through Tn mutagenesis experiments as associated with LPS-5 phage susceptibility are highlighted in red. **D**, Experimental validation of O-antigen/LPS as the receptor for LPS-5. Phage suppression (Suppression ratio) was assessed for clinical *Pa*. A value above the treshold indicates susceptibility to the LPS-5 phage, while values below indicate phage resistance. The suppression ratio was calculated as the area under the curve for the non-phage-treated condition minus the phage-treated condition, divided by the AUC of the non-treated sample. **E**, Colony morphology displaying rugae (right) indicates the presence of LPS with O-antigen (O-AG), whereas smooth colonies (left) lacking rugae suggest mutations in the O-AG within the LPS.

To validate the role of LPS as a receptor in LPS-5 infection, we conducted a suppression assay for ten clinical isolates (Figure 5D). The function of the O-antigen in these clinical isolates was also assessed using a rugae O-AG/LPS assay (Figure 5E). All five bacterial strains resistant to the LPS-5 phage did not form a colony biofilm rugae (rough colony morphotype), suggesting mutations in their O-antigen within the LPS (Supplementary Figure 6). Conversely, 2 out of the 5 susceptible strains exhibited a rugae phenotype, indicative of LPS containing the intact O-antigen.

Consistent with this role for LPS as a receptor for LPS-5, Mutations in LPS biosynthetic genes in these strains were studied alongside with LPS-5 phage infection. The genes *migA* and *wzy* had the most mutations, with some mutations also found in the genes *ssg* and *wapH*. However, due to the limited sample size, no associations between single mutations and susceptibility to phage infection could be identified (Supplementary Table 4).

### The presence of bacterial defense systems is not associated with lytic phage susceptibility

We were interested to observe that bacterial defense systems were not identified in the GWAS, suggesting that these defense systems may not be involved in initial stages of phage infection.

To confirm the potential role of bacterial defense systems in phage infection, we first screened the bacterial genomes of 80 clinical isolates comprising the LPS-5 susceptibility dataset with Padloc (Figure 6A) and identified a total of 123 defense systems (Figure 6B). We then assessed the individual relationship between each defense system and phage susceptibility using a Wilcoxon test. For example, RM type I system was tested as shown in Figure 6C. The results revealed that no defense system was significantly associated with the phenotype of phage susceptibility or resistance (Wilcoxon test, p-value > 0.05 for all) (Figure 6D).

To consider the combined effect of these defense mechanisms on phage susceptibility, we included the defense systems depicted in Figure 6D in a Generalized Linear Model (GLM). None of the systems were significantly associated with susceptibility to either LPS-5 or OMKO1 susceptibility, reaffirming the findings of the Wilcoxon analysis (GLM, p-value > 0.05) (Figure 6E).

These results suggest that phage defense systems do not play a role in determining susceptibility or resistance to lytic phages LPS-5 and OMKO1.

**Figure 6:**
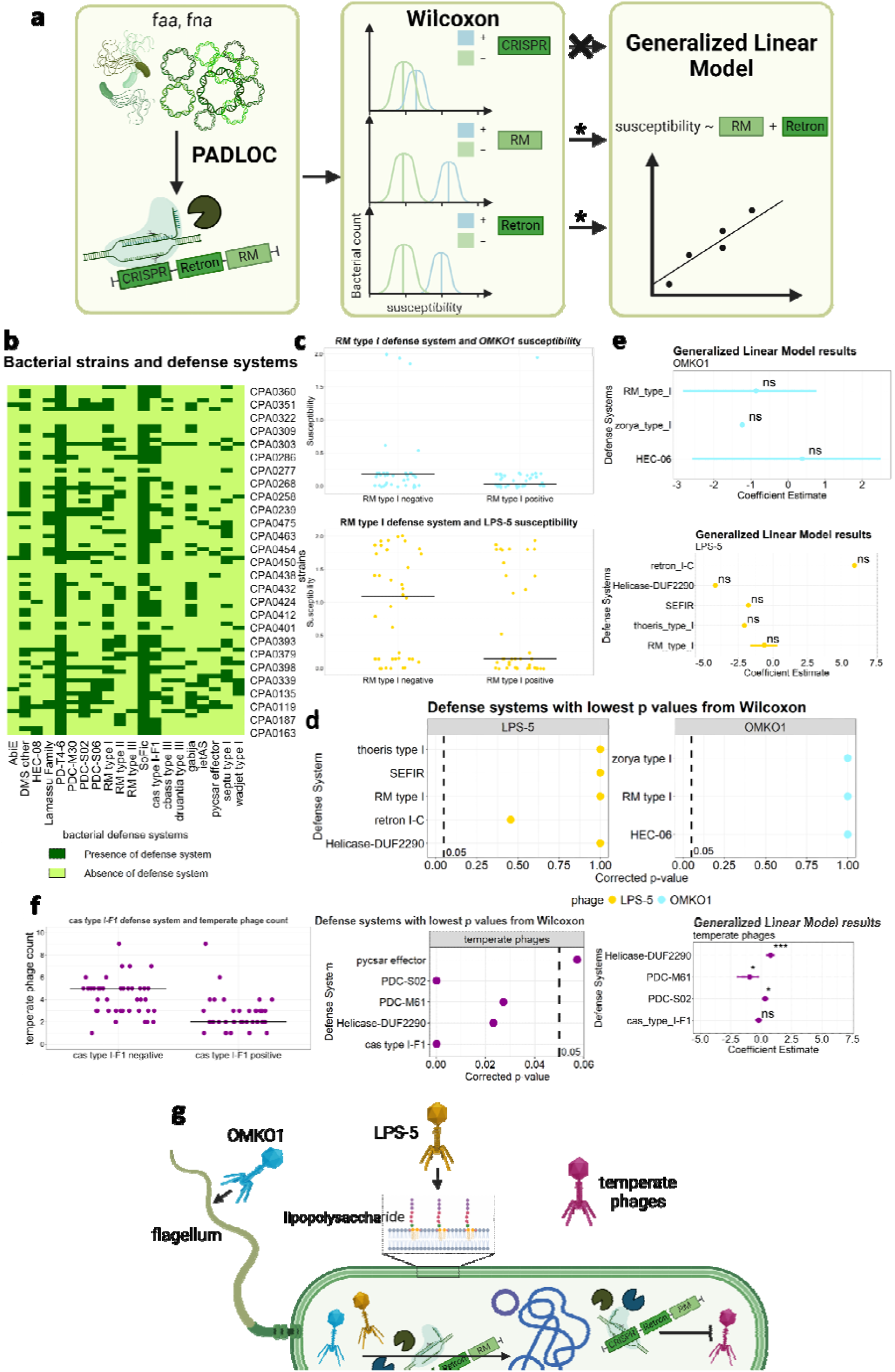
Presence of bacterial defense systems is not associated with lytic phage susceptibility. **A**, Defense System Analysis Workflow. Clinical isolates were screened for defense systems using the Prokaryotic Antiviral Defense LOCator (Padloc). Pre-selection of defense systems was conducted via a Wilcoxon signed-rank test, followed by their inclusion in a generalized linear model to assess potential associations between the presence or absence of these defense mechanisms and phage susceptibility. **B**, Presence or absence of bacterial defense systems across bacterial strains, shown in dark and light green respectively. **C**, Phage susceptibility for isolates with and without RM Type I. Each point represents one bacterial strain grouped into RM type I negative and positive categories. The median is represented by the horizontal line. The first plot illustrates the data for OMKO1, while the second plot includes all data points for LPS-5. **D**, Corrected p-values from the Wilcoxon test for the defense systems with the lowest p-values. **E**, Coefficient estimates for the GLM. The defense systems identified in the Wilcoxon analysis were incorporated into a Generalized Linear Model (GLM). None of the systems are significantly associated with OMKO1 susceptibility or LPS-5 susceptibility. **F**, Bacterial defense systems are associated with the number of temperate phages in the bacterial genome. Similar to the previous analysis, we first investigated the association between the count of temperate phages and the presence of defense systems using a Wilcoxon test, followed by a GLM analysis. **G**, Concluding Schematic. This study suggests that receptors alone determine the host range of lytic phages. We identified the flagellum as a receptor of OMKO1 and lipopolysaccharide as the receptor of LPS-5. Although no significant relationship was found between bacterial defense systems and lytic phage infection, defense mechanisms may play a role in protecting against temperate phages.

### The presence of bacterial defense systems is associated with temperate phage infection

To investigate the association between bacterial defense systems and temperate phage infection, we screened the genomes of clinical isolates for the presence of temperate phages using VIBRANT and BLAST. We first assessed the individual relationship between each system and temperate phages using a Wilcoxon test. The results showed that the presence of 4 defense systems, namely PDC-S02, PDC-M61, Helicase-DUF2290 and cas type I-F1 was significantly associated with the number of temperate phages (Wilcoxon test, p-value < 0.05 for all, Figure 6F). Two of these defense systems (PDC-S02 and Helicase-DUF2290) exhibited a positive correlation with the number of phages, while the other two systems (PDC-M61 and Cas type I-F1) showed a negative correlation.

Next, to consider the combined effect of these defense mechanisms on the number of prophages, we included the significant defense systems in a Generalized Linear Model. We found that three of the aforementioned defense systems were significantly associated with the number of prophages (GLM, p-value < 0.05 for all).

To account for correlations arising from defense systems encoded within temperate phage islands in the bacterial genome, we repeated the analysis after removing all defense systems located within prophage regions. The defense systems Helicase-DUF2290 and PDC-S02 still showed a significant correlation with the number of temperate phages in the bacteria (Supplementary Figure 7).

Together, these results suggest that certain bacterial defense systems may play a role in the regulation of temperate phage infection in clinical isolates.

## Discussion

In this work, we investigated the bacterial genes that influence the host range of two lytic phages, OMKO1 and LPS-5, in a large set of *Pa* clinical isolates collected from pwCF. Our results implicate bacterial structures known to serve as receptors for these and other phages, which is consistent with the observation that the emergence of bacterial resistance to phages is often associated with the loss of receptor structures^19,32,33^.

Our results demonstrate that the phage OMKO1 can utilize the flagellum as a receptor for host infection, which has previously been shown for other phages^34–39^. We speculate that the phage binds to a specific region of the flagellum, serving as a receptor, and then uses the flagellar motion to approach the bacterial surface. The evolution of bacterial resistance against flagellotropic phages is known to involve mutations and the downregulation of flagella^39^. Initial work identified OprM as the receptor for OMKO1^31,34,40^ but other more recent results suggest that other structures like the type IV pilus and flagellum are used by OMKO1^41^. In this context, our findings provide critical evidence supporting the flagellum as the functional receptor for OMKO1 infection.

Regarding the phage LPS-5, our results suggest that it uses the lipopolysaccharide, specifically the O-antigen, as a receptor for infecting its host, similar to many other phages that bind the O-antigen as their receptor^42,43^. This hypothesis is supported by the identification of genes linked to the biosynthesis machinery of the O-antigen in the transposon mutagenesis assay. Our O-antigen experiments further provide support for the role of the O-antigen as the receptor for phage LPS-5^44^. Other genes controlling rugae formation, such as those involved in quorum sensing, may also play a role in influencing rugae formation^45^.

Other pathways enriched in susceptibility-associated genes were identified besides ones contributing to receptor structures. Experiments for both phages showed enrichment of numerous metabolic pathways in the GWAS and Tn assay, including citrate cycle, fatty acid metabolism, carbon metabolism, pyrimidine metabolic processes, amino sugar and nucleotide sugar metabolism and others. These findings are consistent with previous studies that have highlighted the relationship between phage infection and metabolic changes in host bacteria^46–48^. In one study, multi-phage resistance was found to be associated with reduced fitness in a PAO1 population, but not in clinical isolates. The authors hypothesized that clinical strains, having co-evolved with phages, controlled phage infection not solely through direct modification of genes responsible for surface receptor biosynthesis, but rather through global regulation of cell metabolism^46^. Another study has shown that different *Pa* phages can not only deplete metabolites but also completely reorganize the metabolism in a phage- and infection-specific manner, eliciting an increase in pyrimidine and nucleotide sugar metabolism^47^. Thus, phage infection can exert considerable stress on bacteria, leading to alterations in their metabolic profiles.

Pathways for mismatch repair, particularly involving the protein MutL and DNA repair were found to be linked with phage infection. Some viruses employ non-homologous end joining (NHEJ) for viral genome circularization, and the loss of NHEJ could potentially serve as an anti-viral strategy for the host bacteria^49^. Another study also identified mutations in genes unrelated to the corresponding phage receptor molecules but rather involving deletions in the DNA mismatch repair protein MutL^48^.

Pathways associated with biofilm formation and exopolysaccharide biosynthesis were found to be linked with phage infection (Figure 4A). This finding is consistent with a previous study that demonstrated changes in pathways associated with biofilm and exopolysaccharide production in phage-resistant *Pa* strains^48^. Biofilm formation and the production of extracellular polysaccharides are important in bacterial defense and may influence susceptibility to phage infection.

Surprisingly, we did not observe any association between known phage defense systems present in these clinical isolates to susceptibility or resistance to the pair of phages studied. This finding contrasts with previous studies implicating defense mechanisms in protection from phage infection^50–52^ and a recent study reporting an accumulation of defense systems in phage-resistant *Pa* strains^53^. However, our findings are consistent with other studies that have likewise not identified roles for bacterial defense mechanisms in *Pa* resistance to lytic phages^26,54,55^. This discrepancy may be explained by high specificity of phage defense mechanisms to particular phages, which may not offer generalized protection. Alternatively, generalized protection may exist, with the presence of sophisticated phage strategies to counteract bacterial defense systems confounding its observation^25,28^. Notably, a very recent study in *E. coli* reported findings in line with our results^56^. The researchers demonstrated that phage-bacteria interactions in *E. coli* were primarily determined by adsorption factors rather than antiphage defense systems, which played only a marginal role. This independent confirmation in another bacterial species highlights the broader relevance of our findings and suggests that phage defense systems may not be the primary determinants of lytic phage susceptibility in multiple bacterial hosts. To this point, it is known that some “jumbo” phages such as OMKO1 can form a phage nucleus during intracellular replication that can physically protect against DNA-targeting defense systems^57^.

Interestingly, we observe that defense systems are associated with the number of temperate phages integrated in the bacterial genome. Two defense systems, PDC-S02 and Helicase-DUF2290, exhibited a significant positive correlation with the number of phages. Conversely, two other systems, PDC-M61 and Cas type I-F1, showed a negative correlation. Temperate phages may accumulate in a host, making them more suitable targets for adaptive immunity than lytic phages. Defense mechanisms are sometimes encoded in prophages, which could confer a competitive advantage to bacteria harboring temperate phages^58^. Conversely, other defense mechanisms may provide protection against temperate phages^59^. We identified 23 defense systems encoded in 12 prophages in 11 isolates across the 80 isolates from the LPS-5 susceptibility dataset. The positive relationship between some defense systems and number of prophages was still present when omitting prophage-encoded defense systems, suggesting that other factors are driving this relationship. These results are specific to the experimental design and phages studied. Different approaches exploring the relationships and co-evolution between phages and hosts over an extended period, particularly those focusing on the development of phage resistance, may yield different outcomes, as shown by a recent study suggested an accumulation of defense systems in phage-resistant *Pa* strains^53^.

This study is limited by the type of analyses presented here. While Tn mutagenesis assays can identify genes linked to phage susceptibility, they do not reveal genes associated with phage resistance. A transposon insertion into such a gene (loss-of-function) would not confer protection against phage, making it impossible to study a connection with phage infection. Future studies could incorporate gain-of-function assays^60^ or gene cloning for further investigation^55^. In this work, the GWAS may reveal subtle genetic changes associated with both phage susceptibility and phage resistance, making it a complementary tool to Tn mutagenesis assays for identifying the genetic determinants of phage susceptibility.

Padloc represents a robust tool for identifying bacterial defense systems within bacterial strains, being one of the most updated platforms^61^. However, it’s important to acknowledge that the presence of a gene in the genome does not guarantee its expression and functionality, and the tool may not encompass all known defense systems, let alone undiscovered ones. Moreover, defense systems across bacterial species and strains are highly diverse^62^. This diversity, along with their potential interactions or analogies, may reduce the power to detect individual systems and their correlation with phage susceptibility. Nonetheless, Padloc remains the best choice for comprehensive screening, given the impracticality of manual identification.

In conclusion, this study identified genes and structural proteins crucial for phage susceptibility and resistance (Figure 6G). Specifically, flagella were found to play a role in OMKO1 phage infection, while the O-antigen in lipopolysaccharide was identified as important for LPS-5 phage infection. Conversely, our data does not support the widely held view that phage defense mechanisms play a central role in determining phage susceptibility. These results may contribute to our understanding of the constraints of phage infection and could facilitate the development of phage therapy.

## Method and Materials

### Bacteriophages and *Pa*

The phages investigated in this study include the Myoviridae phages LPS-5 and OMKO1. LPS-5 is used in the Cystic Fibrosis Bacteriophage Study at Yale (CYPHY) (NCT04684641) ^16^, while OMKO1 (outer-membrane-porin M knockout dependent phage 1) was isolated at Yale University ^31^ (Table 1, Supplementary Figure 8). PAO1 and the clinical *Pa* isolates used in this study were obtained from our research group (Table 2).

**Table 1:** Bacteriophages.

| Name | Family | Host | Origin |
| --- | --- | --- | --- |
| OMKO1<br>(outer-membrane-porin M knockout dependent phage 1) | Myoviridae | *Pseudomonas aeruginosa* PAO1 | B. K. Chan et al., Yale University<sup>31</sup> |
| LPS-5 | Myoviridae | *Pseudomonas aeruginosa* PAO1 | B. K. Chan et al., Yale University<sup>16</sup> |

**Table 2:** Bacterial strains.

| Strain | Species | Application |
| --- | --- | --- |
| PAO1 | *Pseudomonas aeruginosa* | Tn assay, experimental validation |
| clinical isolates | *Pseudomonas aeruginosa* | lytic plaque assay, swimming assay, suppression assay |

### Transposon mutagenesis assay

The creation of a transposon mutagenesis library was previously described by Larry Gallagher et al. in the Manoil group at the University of Washington.^**63**^ Tn mutagenesis was performed via conjugation of the *Escherichia coli* SM10λpir/pIT2, a Tn5-derived transposon, followed by selection for *Pa* derivatives carrying the transposon tetracycline-resistance element. This results in a different transposon insertion and gene disruption in each mutant strain. The composition of transposon mutant pools was characterized using the Tn-seq circle method (35). The *Pa* library, along with the phages LPS-5 and OMKO1, underwent expansion and titration. For the assay, a bacterial concentration at least 1000 times greater than the number of unique mutants (1×10^5^) was utilized, resulting in an inoculation of 1×10^8^ colony-forming units (cfu) in 25 ml of LB medium. The phages were added at two multiplicities of infection (MOI), 20 and 0.1, corresponding to 2×10^9^ and 1×10^7^, respectively plaque-forming units per milliliter of the medium (pfu/mL). Bacterial cultures were grown to an optical density (OD) of 0.2-0.4, for a duration of 30 minutes to 3 hours depending on the MOI, which is equivalent to 1-3 phage propagation cycles. Subsequently, the bacterial cultures were harvested and prepared for genomic DNA (gDNA) extraction. Transposon region amplification of the gDNA was conducted using the C-tailing method, a PCR reaction with one primer targeting a C-tail region near the 3’ end of the transposon and the second primer complementary to bacterial genomic DNA. Following PCR Tn amplification, DNA was quantified to ensure equal DNA amounts for subsequent Tn-Seq protocol steps. This was followed by library preparation to add adapter sequences necessary for Illumina sequencing. Transposons were sequenced for distinct timepoints and conditions: before phage administration (T0), after killing assay with phage (T1P sequence condition at timepoint 1, see above for durations), and without phage (T1 condition). Reads were mapped to the bacterial genome and quantified. Abundant transposon counts indicated bacterial survival, suggesting that the disruption of the gene played a protective role against phage infection. This allowed the identification of genes crucial for phage infection. Conversely, minimal or absent transposon counts signified bacterial death despite the non-functioning gene and that the disrupted gene was not essential for phage infection.

### Analysis

We first normalized the read counts, dividing the count of a particular gene under a specific condition (timepoint and phage condition) by the sum of all the reads for all the genes at that same timepoint and condition. Following normalization, the gene fitness score was computed by taking the log2 of the normalized gene count at timepoint T1 under the specific phage condition, divided by the count of that gene at timepoint T0 (Equation ( 1 )).

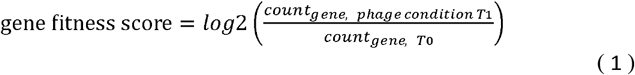

We then performed a Fisher’s test to identify genes significantly enriched in the Tn assay. The Fisher’s exact test calculates the significance of the deviation from the null hypothesis from a contingency table of two categorical variables^64^, in this case phage condition and time.

A functional enrichment analysis was performed using the PANTHER^63^ package to explore these genes further and identify significantly enriched pathways (Table 3). Significant hits were assigned to classes of genes and pathways that were over-represented in the gene set, likely associated with the phage infection phenotype.

**Table 3:** Statistical and analytical tools.

| Tool | Origin | Application |
| --- | --- | --- |
| Fisher's Exact Test for Count Data | R. A. Fisher<sup>64</sup> | Tn assay |
| PANTHER | H. Mi et al.<sup>65</sup> | Functional enrichment analysis |
| Genome-Wide Association Study (GWAS) / De Bruijn Genome-Wide Association Study (DBGWAS) | M. Jaillard et al.<sup>66</sup> | GWAS with lytic plaque assay |
| GraphPad Prism Version 7 | GraphPad Prism | Suppression Ratio calculation |
| Blast | NCBI<sup>67</sup> | Mutations in flagella and LPS associated genes |
| Mafft | Katoh K. et al.<sup>68</sup> | Sequence alignments |
| Cons, EMBOSS | Rice P. et al.<sup>69</sup> | Consensus sequences |
| Prokaryotic Antiviral Defense LOCator (Padloc) | L. J Payne et al.<sup>61</sup> | Bacterial defense systems |
| Wilcoxon signed-rank test | Kurt Hornik<sup>70</sup> | Bacterial defense systems, single system analysis |
| Generalized Linear Model (glm) | Simon Davies<sup>71</sup> | Bacterial defense systems, all systems |
| R Version 4.3 and RStudio | R and RStudio | Analysis, graphs |
| BioRender | BioRender<sup>72</sup> | Schematics |

### Suppression Assay

Cultures from frozen stock were initially incubated overnight at 37°C with shaking at 250 rpm. Following this, overnight cultures of bacterial strains were diluted 1:200 with LB medium (Table 4) and regrown for 2-3 hours until reaching an OD600 of approximately 0.1-0.2. Subsequently, this culture was diluted 1:10 into the wells of a 96-well plate, and phage was added to achieve a MOI of 100, resulting in a total volume of 150 μl per well. The 96-well plate was then incubated at 37°C with shaking for 30 hours, during which the OD600 was measured every 20 minutes using an automated spectrophotometer (Biotek microplate reader). All growth curves were adjusted by subtracting the background curve (LB medium with phage buffer at a concentration of 1 mM MgSO4, 4 mM CaCl2, 50 mM Tris-HCl (pH=7.8), 6 g/L NaCl, and 1 g/L gelatin). The Suppression Ratio, expressed as a percentage, was calculated using GraphPad Prism Version 7.0 software. This ratio was defined as the area under the curve (AUC) for the non-phage-treated condition minus the AUC for the phage-treated condition, divided by the AUC of the non-treated condition.

**Table 4:**
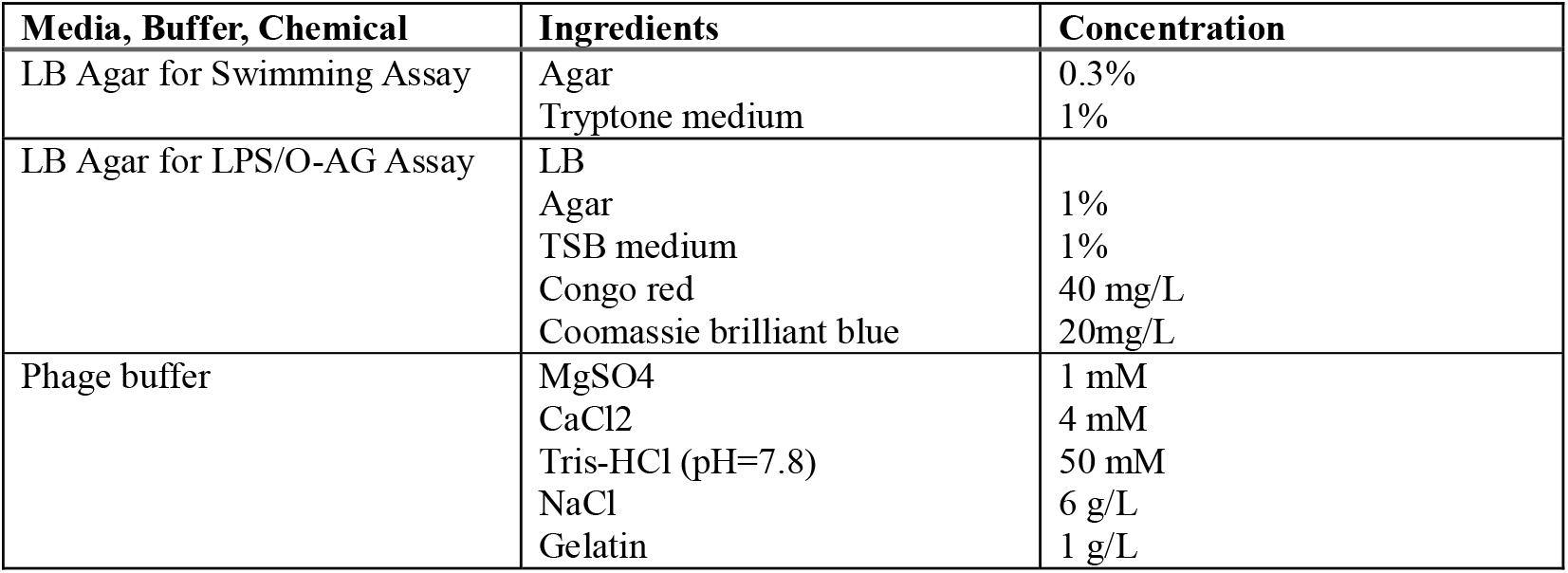
Media, Buffers, Chemicals.

| Media, Buffer, Chemical | Ingredients | Concentration |
| --- | --- | --- |
| LB Agar for Swimming Assay | Agar<br>Tryptone medium | 0.3%<br>1% |
| LB Agar for LPS/O-AG Assay | LB<br>Agar<br>TSB medium<br>Congo red<br>Coomassie brilliant blue | 1%<br>1%<br>40 mg/L<br>20mg/L |
| Phage buffer | MgSO4<br>CaCl2<br>Tris-HCl (pH=7.8)<br>NaCl<br>Gelatin | 1 mM<br>4 mM<br>50 mM<br>6 g/L<br>1 g/L |

### Swimming assay

Following a standardized protocol^**73**^, overnight cultures were initially grown and subsequently diluted before regrowth to an OD of 0.1-0.2. The bacterial suspension was then transferred onto agar plates via puncturing with a pipette tip. These agar plates were prepared using 1% tryptone medium supplemented with 0.3% agar. After incubation for 24 hours at 37°C, the growth zones were measured. The experiment was conducted with a minimum of 5 repetitions.

### LPS/O-AG assay

Following an established protocol^**45**^, 1μl of overnight culture of clinical isolates were spotted onto agar plates containing 1% TSB medium fortified with 40 mg/L Congo red and 20 mg/L Coomassie brilliant blue dyes and solidified with 1% agar. Colony biofilms were grown for 4 days at 25°C before images were taken and the morphology analysed (Supplementary Figure 6).

### Whole genome sequencing and assembly

We collected Pa isolates from sputum samples from individuals with CF and banked them with patient consent under IRB #11197, approved by the Stanford University Institutional Review Board. DNA was extracted using the DNA Easy Kit (Qiagen, 69504, Hilden, Germany) and sequenced with Illumina. We trimmed raw reads using trimmomatic^**74**^ and then assembled them with SPAdes using –isolate and --cov-cutoff auto^**75**^.

### Mutant calling

Flagella genes and LPS biosynthetic gene sequences (Supplementary Table 5) were downloaded from the Pseudomonas genome database^**76**^. These sequences were blasted against the genomes of the clinical isolates using the ncbi-blast software^**67**^. Hits were then aligned using mafft^**68**^. To identify non-synonymous mutations, aligned sequences over a specific length (>20% of the total sequence, typically around >400 bp) were translated. Subsequently, each amino acid was compared to the reference sequence used for the blast analysis. Changes in amino acids were documented, including the amino acid position and the specific alteration. For sequences exhibiting high diversity (sequence similarity < 90%), consensus sequences for both resistant and susceptible strains were generated. Consensus sequences for each variant were derived using the cons function from the European Molecular Biology Open Software Suite (EMBOSS^**69**^).

### Lytic plaque assays

Clinical Pa strains were plated on rectangular LB agar plates, followed by the addition of drops of distinct phages in different dilutions (dilution factors: 0, 1, 3, 5, 7). Each experiment was repeated three times and the phages tested were OMKO1 and LPS-5. The plates were then incubated to allow phage infection and plaque formation.

### Analysis

The assessment of plaques involved several parameters (Equation ( 2 )), including single plaque status (SPS), plaque score of the sample (PS) and reference (RS), and inhibition score (IS). SPS was assigned a value of 1 if single plaque resolution was observed at any phage dilution, and 0 otherwise. When SPS was 0, PS was also assigned 0, however, if SPS was 1, its value was determined by the dilution factor with single plaque resolution. RS represented the dilution factor of the reference strain when SPS was 1, assuming a nonzero plaque score for the reference strain. In cases where SPS was 0, IS was set to the highest dilution factor with signs of bacterial growth inhibition. These parameters collectively contributed to the derived score for each clinical isolate, ranging from 0 to 2. Scores between 0 and 1 indicate phage resistance or inhibition of bacterial growth, while scores between 1 and 2 indicate phage susceptibility. This phenotypic data, combined with the genome sequences of the clinical isolates as genotypic data, was used in the GWAS to identify genes associated with either phage susceptibility or resistance.

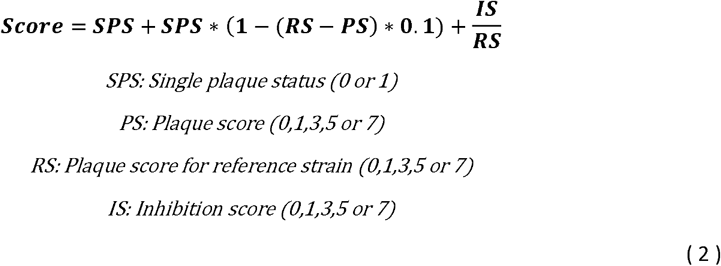

### GWAS

The lytic plaque assay determined which clinical Pa strains were susceptible to the phages OMKO1 and LPS-5. This phenotypic data, combined with the whole-genome sequencing of the clinical isolates, was used in a Genome-Wide Association Study (GWAS) to identify genes associated with either phage susceptibility or resistance.

We performed a De Bruijn Genome-Wide Association Study (DBGWAS). The analysis used a binary phenotype (susceptible vs. resistant) as input together with assembled whole-genome sequences for pangenome analysis. An annotated reference genome was included as well as a phylogenetic tree to account for population structure. DBGWAS uses a compacted de Bruijn graph (cDBG) structure to represent sequence variability across all input bacterial genome assemblies. Nodes in the cDBG were tested for association with the phenotype using a linear mixed model and significant nodes were mapped to the annotated reference genome, yielding phenotype-associated genetic events. Diagnostic analysis was performed using QQ-plots generated with the qqman package.

Given the numerous genes identified by the GWAS in association with phage infection, a functional enrichment analysis was conducted to identify enriched pathways using the PANTHER package. Significant hits are assigned to classes of genes and entire pathways that are over-represented in the gene set associated with the phage infection phenotype.

### Bacterial defense systems and lytic phage infection

The genomes of the clinical isolates were screened for bacterial defense systems using the Prokaryotic Antiviral Defense LOCator (Padloc).^61^ We assessed the association of each system with lytic scores using a Generalized Linear Model (GLM), which was only fitted to systems that were significantly associated with phage resistance or susceptibility, as determined by a Wilcoxon signed-rank test. ^77^ P-values from the Wilcoxon test were adjusted for multiple testing using a Bonferroni correction (each p-value was multiplied by the total number of tests conducted, corresponding to the number of defense systems evaluated). Any corrected p-value exceeding 1 was set to 1. Since no defense systems showed significance in the Wilcoxon test, the systems that exhibited significance prior to p-value correction were selected for inclusion in the GLM ^71^. The GLM was fitted with a binomial distribution, and the AIC criterion ^78^ was employed to compare the fit of different distributions and determine the optimal number of defense systems to include. Additionally, residual diagnostics were performed using the DHARMa package ^79^ to confirm the goodness of fit of the model.

### Bacterial defense systems and temperate phages

We followed the steps described above using the number of temperate phages in the bacterial genome as a response variable. We identified temperate phages using VIBRANT and blast and looked for bacterial defense systems using Padloc. The relationship between each defense system and the number of temperate phages was individually analysed using a Wilcoxon test. Subsequently, significant predictors (p<0.05) were included in a generalized linear model with a Poisson distribution to evaluate the association between these defense mechanisms and the number of temperate phages.

## Data and code availability

Raw reads for bacterial isolates can be accessed through NCBI SRA under BioProject accession PRJNA1188603. All detailed codes used for analysis are available upon request.

## Acknowledgments

We extend our heartfelt gratitude to all members of the Bollyky Laboratory for insightful discussions and support. JDA and PET acknowledge funding support from a research grant award by Howard Hughes Medical Institute. P.L.B. discloses support from National Institutes of Health grants P01AI1960471, R01HL148184-01, K24 AI166718, R21AI194044, and 1R01AI182349-01A1, the Stanford Woods Institute for the Environment, the Stanford-Coulter Translational Research Program, Bio-X, Stanford SPARK, and the Stanford Innovative Medicines Accelerator.

## Authors and Contributors

P.L.B., J.D.P. and D.M.M. conceptualized the main idea of this research project. D.M.M. performed the majority of the experiments and analysis. J.D.P. assembled and processed the bacterial genomes and identified temperate phages. P.L.B., L.A.G and J.D.P. provided feedback on the results. D.M.M. wrote the paper. P.L.B., L.A.G, J.D.P., M.K.K and D.M.M. contributed to the revision of the paper. M.K.K and J.D.A. supported the experimental validation and M. H. provided the TEM pictures. D.M.M. and B.T. created the LPS schematic. E.B.B. collected and banked the clinical isolates and contributed to revisions of the manuscript. R.M. (Co-Founder & CEO of Felix Biotechnology) performed the Transposon mutagenesis experiment and lytic plaque assay.

