## Supplemental Figures and Tables for "Bacterial Receptors but Not Anti-Phage Defense Mechanisms Determine Host Range for a Pair of *Pseudomonas aeruginosa* Lytic Phages"

**Supplemental Tables and Figures**

**Supplementary Table 1: Dataset of clinical isolate strains used for GWAS of the phages OMKO1 and LPS-5.**

|  | OMKO1 dataset | LPS-5 dataset |
| --- | --- | --- |
| 1 | sFB1326 | sFB1326 |
| 2 | sFB1327 | sFB1327 |
| 3 | sFB1344 | sFB1344 |
| 4 | sFB1345 | sFB1345 |
| 5 | sFB1348 | sFB1348 |
| 6 | sFB1366 | sFB1366 |
| 7 | sFB1378 | sFB1378 |
| 8 | sFB1379 | sFB1379 |
| 9 | sFB1389 | sFB1389 |
| 10 | sFB1394 | sFB1394 |
| 11 | sFB1400 | sFB1400 |
| 12 | sFB1401 | sFB1401 |
| 13 | sFB1402 | sFB1402 |
| 14 | sFB1403 | sFB1403 |
| 15 | sFB1404 | sFB1404 |
| 16 | sFB1405 | sFB1405 |
| 17 | sFB1406 | sFB1406 |
| 18 | sFB1407 | sFB1407 |
| 19 | sFB1408 | sFB1408 |
| 20 | sFB1409 | sFB1409 |
| 21 | sFB1410 | sFB1410 |
| 22 | sFB1411 | sFB1411 |
| 23 | sFB1413 | sFB1413 |
| 24 | sFB1415 | sFB1415 |
| 25 | sFB1416 | sFB1416 |
| 26 | sFB1417 | sFB1417 |
| 27 | sFB1418 | sFB1418 |
| 28 | sFB1420 | sFB1420 |
| 29 | sFB1421 | sFB1421 |
| 30 | sFB1422 | sFB1422 |
| 31 | sFB1423 | sFB1423 |
| 32 | sFB1425 | sFB1425 |
| 33 | sFB1426 | sFB1426 |
| 34 | sFB1427 | sFB1427 |
| 35 | sFB1428 | sFB1428 |
| 36 | sFB1429 | sFB1429 |
| 37 | sFB1431 | sFB1431 |
| 38 | sFB1432 | sFB1432 |
| 39 | sFB1433 | sFB1433 |
| 40 | sFB1434 | sFB1434 |
| 41 | sFB1435 | sFB1435 |
| 42 | sFB1436 | sFB1436 |
| 43 | sFB1437 | sFB1437 |
| 44 | sFB1438 | sFB1438 |
| 45 | sFB1439 | sFB1439 |
| 46 | sFB1441 | sFB1441 |
| 47 | sFB1442 | sFB1442 |
| 48 | sFB1443 | sFB1443 |
| 49 | sFB1444 | sFB1444 |
| 50 | sFB1448 | sFB1448 |
| 51 | sFB1451 | sFB1451 |
| 52 | sFB1453 | sFB1453 |
| 53 | sFB1454 | sFB1454 |
| 54 | sFB1455 | sFB1455 |
| 55 | sFB1456 | sFB1456 |
| 56 | sFB1457 | sFB1457 |
| 57 | sFB1458 | sFB1458 |
| 58 | sFB1459 | sFB1459 |
| 59 | sFB1461 | sFB1461 |
| 60 | sFB1462 | sFB1462 |
| 61 | sFB1463 | sFB1463 |
| 62 | sFB1466 | sFB1466 |
| 63 | sFB1467 | sFB1467 |
| 64 | sFB1468 | sFB1468 |
| 65 | sFB1469 | sFB1469 |
| 66 | sFB1470 | sFB1470 |
| 67 | sFB1471 | sFB1471 |
| 68 | sFB1472 | sFB1472 |
| 69 | sFB1473 | sFB1473 |
| 70 | sFB1474 | sFB1474 |
| 71 | sFB1476 | sFB1476 |
| 72 | sFB1477 | sFB1477 |
| 73 | sFB1478 | sFB1478 |
| 74 | CPA0004 | sFB1479 |
| 75 | CPA0027 | sFB1482 |
| 76 | CPA0033 | sFB1483 |
| 77 | CPA0050 | sFB1485 |
| 78 | CPA0061 | sFB1486 |
| 79 | CPA0086 | sFB1487 |
| 80 | CPA0106 | sFB1489 |

**Supplementary Table 2: Consensus sequences**

| Start codon | M |
| --- | --- |
| Stop codon | * |
| No base/sequence identified in blast | **-** |
| *FliC* consensus sequence of resistant strains  MALTVNTNIASLNTQRNLNASSNDLNTSLQRLTTGYRINSAKDDAAGLQISNRLSNQISGLNVATRNANDGISLAQTAEGALQQSTNILQRIRDLALQSANGSNSDADRAALQKEVAAQQAELTRISDTTTFGGRKLLDGSFGTTSFQVGSNAYETIDISLQNASASAIGSYQVGSNGAGTVASVAGTATASGIASGTVNLVGGGQVKNIAIAAGDSAKAIAEKMDGAIPNLSARARTVFTADVSGVTGGSLNFDVTVGSNTVSLAGVTSTQDLADQLNSNSSKLGITASINDKGVLTITSATGENVKFGAQTGTATAGQVAVKVQGSDGKFEAAAKNVVAAGTAATTTIVTGYVQLNSPTAYSVSGTGTQASQVFGNASAAQKSSVASVDISTADGAQNAIAVVDNALAAIDAQRADLGAVQNRFKNTIDNLTNISENATNARSRIKDTDFAAETAALSKNQVLQQAGTAILAQANQLPQAVLSLLR* | |
| *FliC* consensus sequence of susceptible strains  MALTVNTNIASLNTQRNLNNSSASLNTSLQRLSTGSRINSAKDDAAGLQIANRLTSQVNGLNVATKNANDGISLAQTAEGALQQSTNILQRMRDLSLQSANGSNSDSERTALNGEVKQLQKELDRISNTTTFGGRKLLDGSFGVASFQVG---------------------------------------------------------------------------------------------------------------------------------------------------------------------------------------------------------------------------------------------------------------------IDAQRADLGAVQNRFDNTINNLKNIGENVSAARGRIEDTDFAAETANLTKNQVLQQAGTAILAQANQLPQSVLSLLR* | |
| *FliD* consensus sequence of resistant strains  MAGISIGVGSTDYTDLVNKMVNLEGAAKTNQLATLEKTTTTRLTALGQFKSAISAFQTALTALNSNAVFMARTAKSSNEDILKASATQSAVAGTYQIQVNSLATSSKIALQAIADPANAKFNSGTLNISVGDTKLPAITVDSSNNTLAGMRDAINQAGKEAGVSATIITDNSGSRLVLSSTKTGDGKDIKVEVSDDGSGGNTSLSQLAFDPATAPKLSDGAAAGYVTKAANGEITVDGLKRSIASNSVSDVIDGVSFDVKAVTEAGKPITLTVSRDDAGVKDNVKKFVEAYNTLTKFINEQTVVTKVGEDKNPVTGALLGDASVRALVNTMRSELIASNENGSVRNLAALGITTTKDGTLEIDEKKLDKAISADFEGVASYFTGDTGLAKRLGDKMKPYTDAQGILDQRTTTLQKTLSNVDTQKADLAKRLAALQEKLTTQFNLLSAMQDEMTKRQKSITDNLASLPYGSGKKT* | |
| *FliD* consensus sequence of susceptible strains  No blast sequence identified | |
| *FlgK* consensus sequence of resistant strains  MSDLLSIGLSGLGTSQTWLTITGHNITNVKTPGYSRQDAIQQT--PQFSGAGYMGSGSQIVDVRRLASDFLTGQLRNATSQNSELSAFLGQIDQLNSLLADNTTGVSPAMQRFFSALQTAAQNPSSTEAREAVLAQAQGLSKTFNTLYDQLDKQNSLINQQLGALTSQVNNLSQSVAEYNDAIAKAKSAGAVPNDLLDARDEAVRKLSEMVGVTAVTQDDNSVSLFIGSGQPLVVGNTVSTLSVVPGLDDPTRYQVQLTLGDSTQNVTRLVSGGQMGGLLAYRDTVLDSSYNKLGQLALTFADTVNKQLGQGLDLAGKAGANLFGDINDPDITALRVLAKNGNTGNVHANLNITDTSKLNSSDFRLDFDGTNFTARRLGDDASMQVTVSGTGPYTLSFKDANGVDQGFSVTLDQLPAAGDRFTLQPTRRGASDIETTLKNASQLAFAGSARAEATTNNRGSGAIGQPNLVGGPSPIDPAVLQNAFGANGLPLSATVSADGKTYTMTSPLPAGWSYVDKDGNALPGSPTLNSGTSNSVRMAYTDPGSGQTYTYEFNLSNVPQTGDSFTLSFNKDGISDNRNALNLNALQTKPTVGGTGSTGSTYNDAYGGLVERVGTLTAQARASADASQTVLKQAQDSRDSLSGVSLDEEAANLIQFQQYYSASAQVIQVARSLFDTLIGAFR* | |
| *FlgK* consensus sequence of susceptible strains  MSDLLSIGLSGLGTSQTWLTITGHNITNVKTPGYSRQDAIQQTQVPQFSGAGYMGSGSQIVDVRRLASDFLTGQLRNATSQNSELSAFRSQIEQLDGLLSNTTTGVSPAMQRFFAALQAAANNPSSTEAREAVLAQAEGLGKTFNTLYDQLDKQNSLINQQLGALASQVNHLSQSVASYNDAIAKAKSAGAVPNDLMDARDEAVRKLSEMIGVTAVTQDDNSVSLFIGSGQPLVVGNTVSTLSVVPGLDDPTRYQVQLSNGNSIQNVTGLVSGGEMGGLLAYRNSALDSSYNKLGQLAITLADTINKQLGQGLDLAGKAGANLFGDINDPDITALRVLAKNGNTGNVHANLNITDTSKLNSSDFRLDFDGTSFTARRLGDDASMQVTVSGTGPYTLSFKDANGVDQGFNLTLDQLPAAGDRFTLQPTRRGAADIEATLKNASQLAFAGTARTESTTENRGTGKIGAPTLTSGPSPVDPTVLQGAFGPNGISLGATLSADGKTYTLSSPLPAGWSYVDKDGNALTGSPTLTSGNTNTVRMAFTDP-SGQKYSYEFELSGVPQNSDSFKLGFNDKGISDNRNALNLLALQTKPTVGGTDNTGSTYNEAYGGLVERVGTLTAQVRASSEASATVLKQAQDSRDSLSGVSLDEEAANLIQFQQYYGASAQVIQVARTLFDTLIGAFR* | |
| *FlgL* consensus sequence of resistant strains  MRISTIQAFNNSVSGISRNYADLTRTQAEISAGKRLLTPADDPVGAVRLLQLNQEQALNSQYKSGITAAKNSLQQEETILNSVGTVIHRIREIAVQAGNGGLDASDKNALATELAQREDELLNLLNSRDASGKYLFSGSQGDTQPFVRNPDGTYSYNGDEGQREVQIASSTFIAISDNGKILFESGSNANRVSTGKDAAGLDASGNPSDSSISLGLVTDKEAYDTVFPSSTPPLASDGVGIHFTDAKNYVVYDLKTIPPGYDWSTSDPNPP-SFATLASGKIDDNAQTSDFITFGGVKVQIDGEPQGGDTFSIKRQPDQEKRSLLNTVSDLRKALLSAEDTPAGNLAIRDAVGVAISNLDSSNSQILTGQGRIGARMNVAESTETFIDDVTLVNTAVISQIQDLDYPEALSRLTLQSTIMDAAQQSFVKIRGLSLFNYLS* | |
| *FlgL* consensus sequence of susceptible strains  MRISTIQAFNNSVNGISRNYADLNRTFEQISTGKRILTPADDPVGSVRLLRLDQEQGLNEQYKTGMTEAKNSLSQEETILRSVGNVLQRIREIAGQAGDGALDSNDKKSLASELRQREDELLNLLNSRDASGKYLFSGSQGSVQPFVRNEDGTYSYMGDESQREVQIASSTRIPVSDSGKVLFEDIVNAARLDT-------AAAGNTGDGRISVGLVEDELAFDSQFPASNPPAATDGFNIHFVSDKEYVVYDPKSLPPGYDWTTYDPNSPPAWQ-LSKGAIDDDPKTIDKVLYAGVSVTIDGTPKAGDEFNVNYKPGSEKRSLLNVVSDLRKALESSTDNQAGNDAIRDATAVALTNLSAVAAAVDGGQGKIGARLNTVESTETFIDDVKLVNASVMSQIQDLDYAEALSRLSLQSTIMDAAQQSYVKIQGLSLFNYL-- | |

**Supplementary Table 3: Various bacterial sequences and mutations were identified in isolates resistant to OMKO1.** The biggest differences were observed in the genes *fliC*, *fliD*, *flgK*, and *flgL*. Sequence similarity for these genes within resistant and susceptible strains was clustered, and consensus sequences for both resistant and susceptible strains were generated (Supplementary Table 2). Single mutations were identified in other flagella genes.

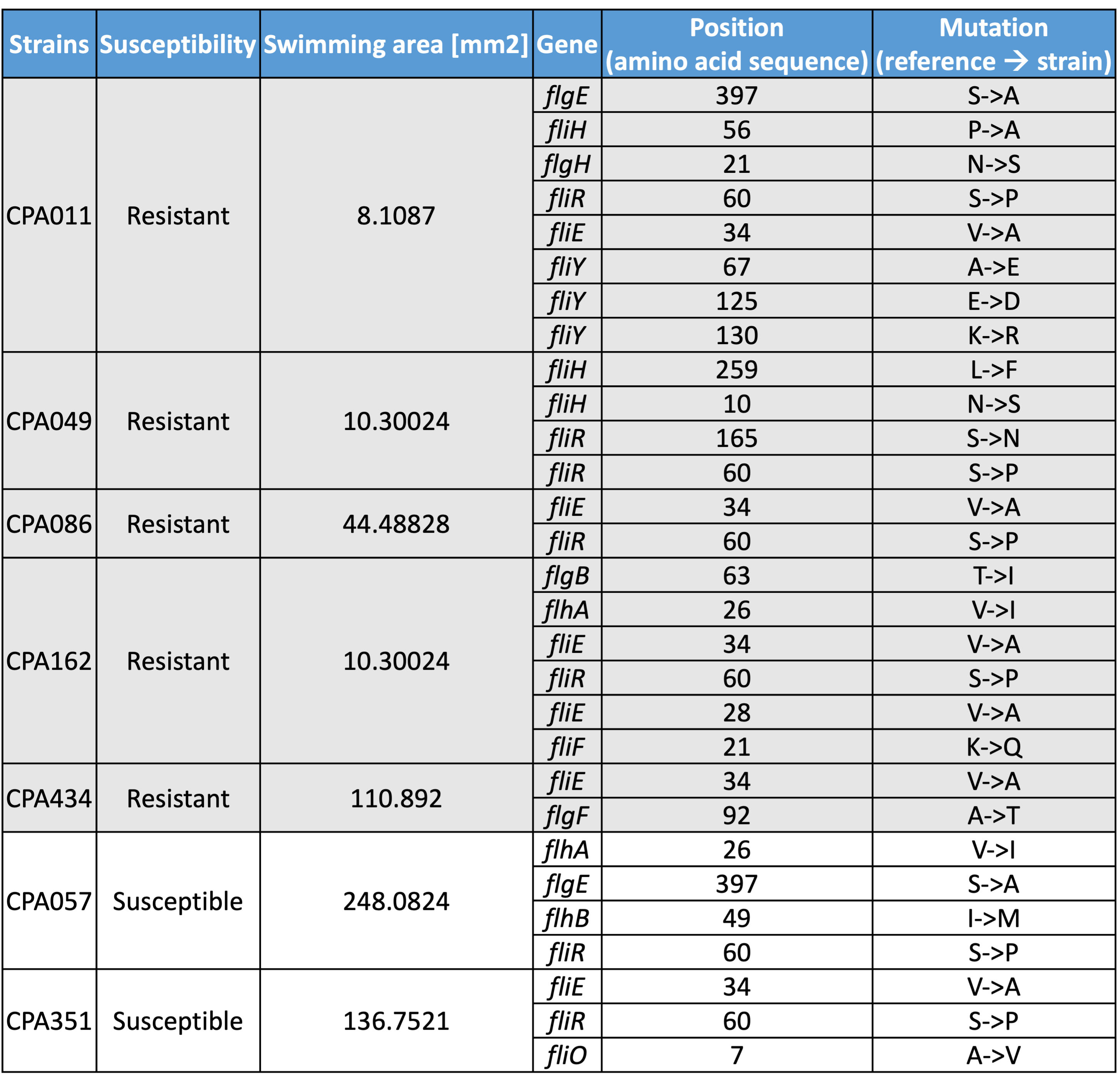

**Supplementary Table 4: Mutations identified in LPS biosynthetic genes.** Most mutations were observed in the genes *migA* and *wzy*, with some mutations also found in the genes *ssg* and *wapH*.

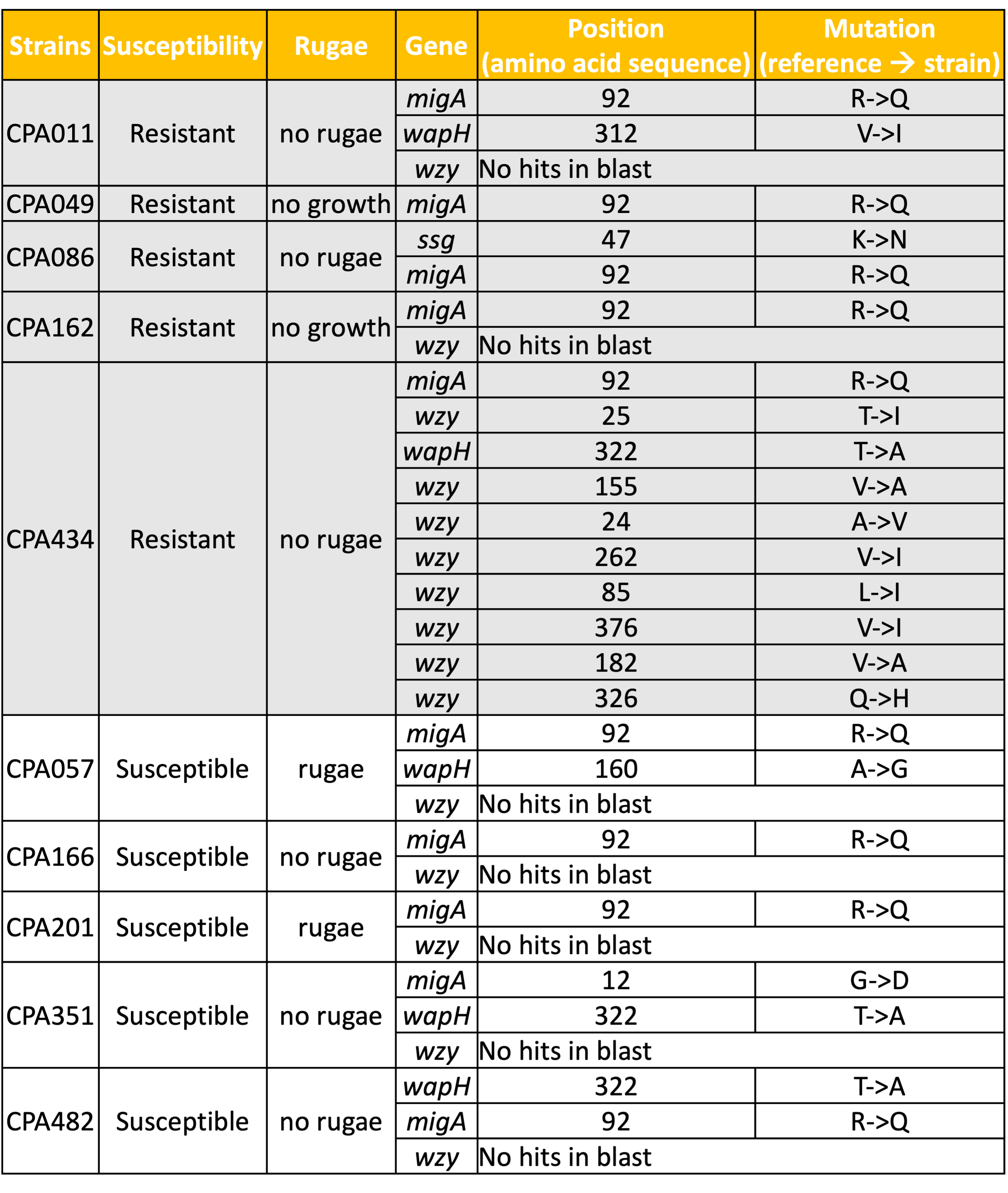

**Supplementary Table 5: Flagella and LPS biosynthetic reference sequences from Pseudomonas genome database.**^76^ Flagella genes and LPS biosynthetic gene sequences were downloaded from the Pseudomonas genome database.

| **Flagella gene** | **RefSeq** |
| --- | --- |
| *fliD* | [NP_249785.1](http://www.ncbi.nlm.nih.gov/protein/NP_249785.1) |
| *fliC* | [NP_249783.1](http://www.ncbi.nlm.nih.gov/protein/NP_249783.1) |
| *flgL* | [NP_249778.1](http://www.ncbi.nlm.nih.gov/protein/NP_249778.1) |
| *flgK* | [NP_249777.1](http://www.ncbi.nlm.nih.gov/protein/NP_249777.1) |
| *flgE* | [NP_249771.1](http://www.ncbi.nlm.nih.gov/protein/NP_249771.1) |
| *flgG* | [NP_249773.1](http://www.ncbi.nlm.nih.gov/protein/NP_249773.1) |
| *flgH* | [NP_249774.1](http://www.ncbi.nlm.nih.gov/protein/NP_249774.1) |
| *flgI* | [NP_249775.1](http://www.ncbi.nlm.nih.gov/protein/NP_249775.1) |
| *flgB* | [NP_249768.1](http://www.ncbi.nlm.nih.gov/protein/NP_249768.1) |
| *flgC* | [NP_249769.1](http://www.ncbi.nlm.nih.gov/protein/NP_249769.1) |
| *flgF* | [NP_249772.1](http://www.ncbi.nlm.nih.gov/protein/NP_249772.1) |
| *fliE* | [NP_249791.1](http://www.ncbi.nlm.nih.gov/protein/NP_249791.1) |
| *fliO* | [NP_250136.1](http://www.ncbi.nlm.nih.gov/protein/NP_250136.1) |
| *fliP* | [NP_250137.1](http://www.ncbi.nlm.nih.gov/protein/NP_250137.1) |
| *fliQ* | [NP_250138.1](http://www.ncbi.nlm.nih.gov/protein/NP_250138.1) |
| *fliR* | [NP_250139.1](http://www.ncbi.nlm.nih.gov/protein/NP_250139.1) |
| *fliF* | [NP_249792.1](http://www.ncbi.nlm.nih.gov/protein/NP_249792.1) |
| *fliG* | [NP_249793.1](http://www.ncbi.nlm.nih.gov/protein/NP_249793.1) |
| *fliY* | [NP_249005.1](http://www.ncbi.nlm.nih.gov/protein/NP_249005.1) |
| *fliM* | [NP_250134.1](http://www.ncbi.nlm.nih.gov/protein/NP_250134.1) |
| *fliN* | [NP_250135.1](http://www.ncbi.nlm.nih.gov/protein/NP_250135.1) |
| *flhA* | [NP_250143.1](http://www.ncbi.nlm.nih.gov/protein/NP_250143.1) |
| *flhB* | [NP_250140.1](http://www.ncbi.nlm.nih.gov/protein/NP_250140.1) |
| *fliH* | [NP_249794.1](http://www.ncbi.nlm.nih.gov/protein/NP_249794.1) |
| *fliI* | [NP_249795.1](http://www.ncbi.nlm.nih.gov/protein/NP_249795.1) |
| *fliJ* | [NP_249796.1](http://www.ncbi.nlm.nih.gov/protein/NP_249796.1) |
| **LPS biosynthetic gene** | **RefSeq** |
| *migA* | [NP_249396.1](http://www.ncbi.nlm.nih.gov/protein/NP_249396.1) |
| *wzy* | [NP_251844.1](http://www.ncbi.nlm.nih.gov/protein/NP_251844.1) |
| *wapH* | [NP_253691.1](http://www.ncbi.nlm.nih.gov/protein/NP_253691.1) |
| *ssg* | [NP_253688.1](http://www.ncbi.nlm.nih.gov/protein/NP_253688.1) |

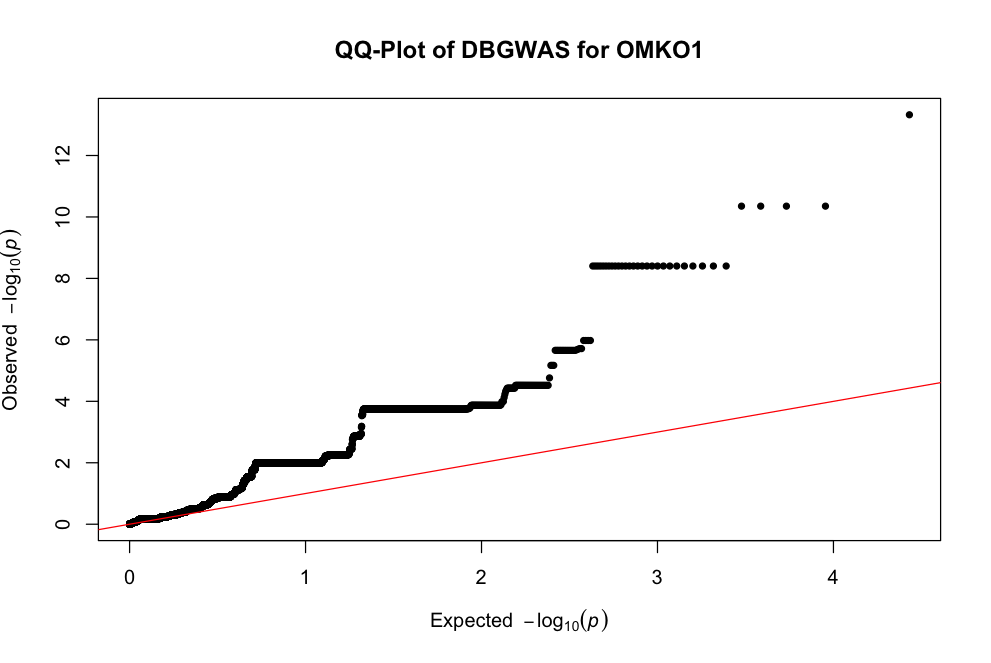

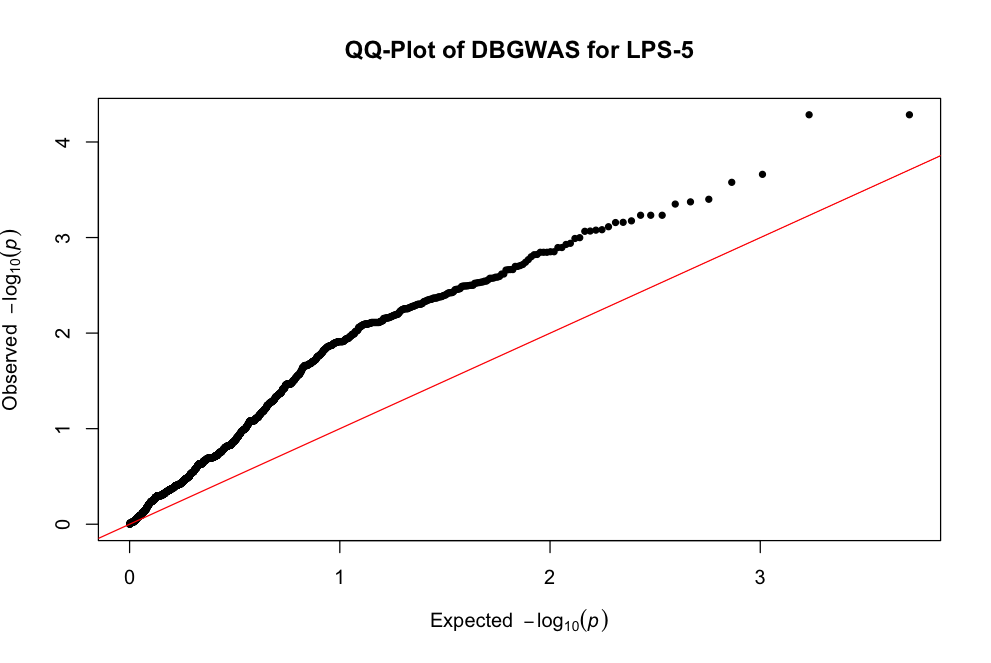

**Supplementary Figure 1: QQ-plots of DBGWAS analyses for phages OMKO1 and LPS-5.** The expected distribution is shown with the red line. Deviations indicate remaining population structure or other confounding variables.

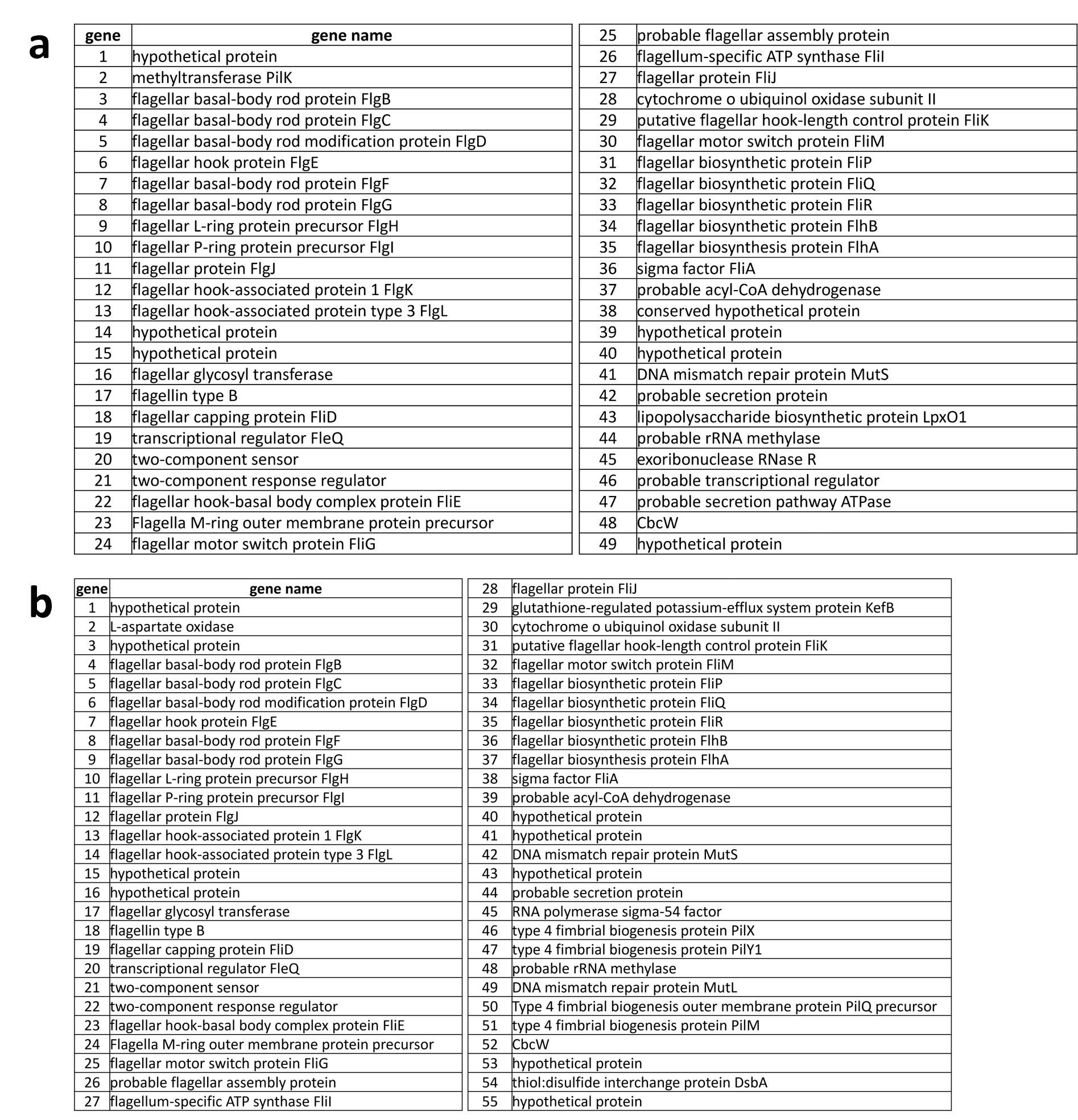

**Supplementary Figure 2: Flagella-Associated Genes are required for OMKO1 Phage Infection.** Full legend to the volcano plots in Figure 2B and Supplementary Figure 3A for OMKO1 at MOI 0.1 (a.) and MOI20 (b.). Volcano plot describes genes with a significant difference in counts (fold change) between conditions with and without OMKO1 phage.

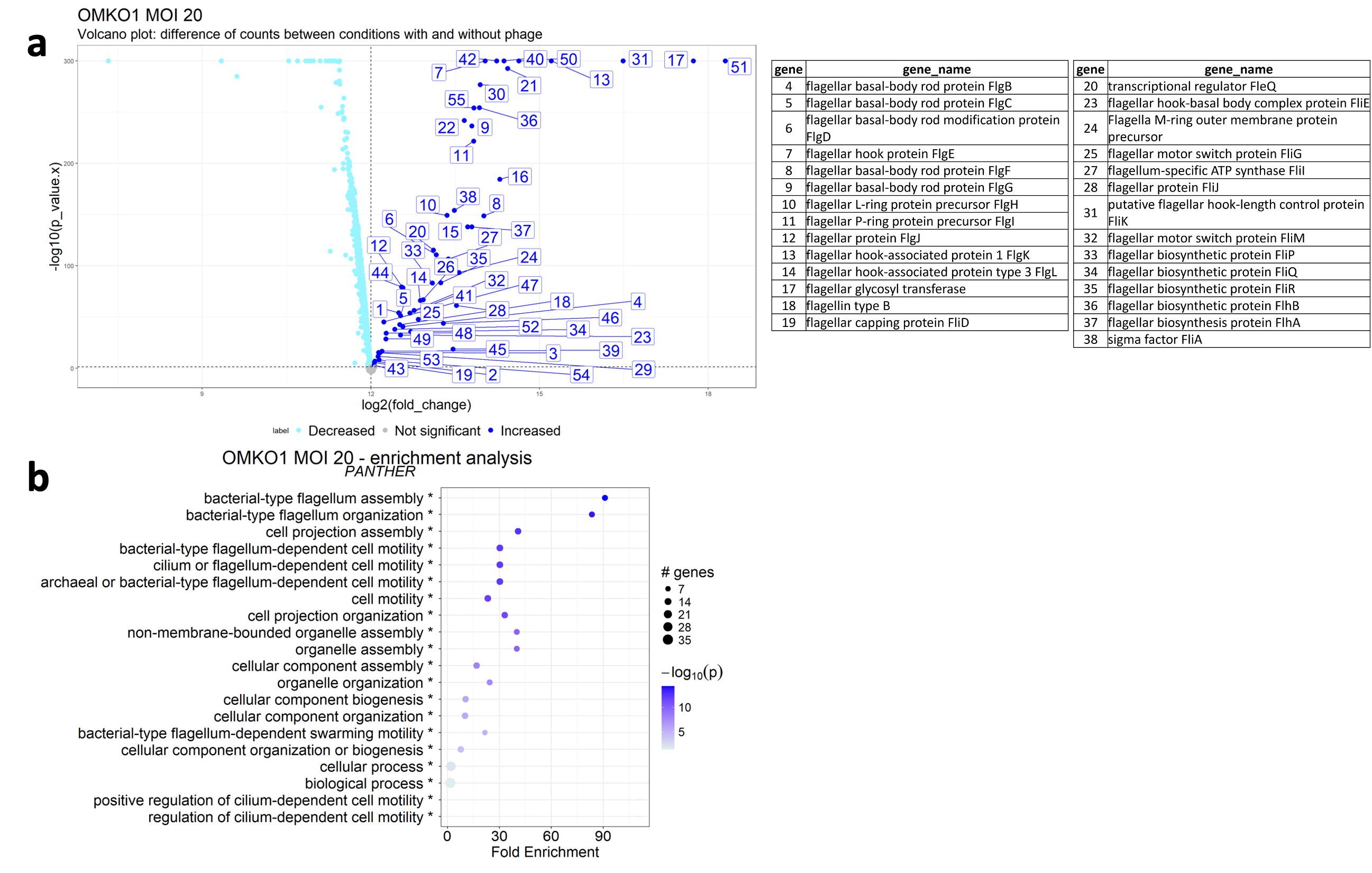

**Supplementary Figure 3:** **Flagella-Associated Genes are required for OMKO1 Phage Infection A,** This volcano plot describes genes with a significant difference in counts (fold change) between conditions with and without OMKO1 phage at MOI 20. Dark blue signifies a significant increase in gene counts, bright blue stands for a significant decrease, and gray represents non-significant genes. **B,** Functional enrichment analysis results for the OMKO1 phage at MOI 20 with PANTHER.

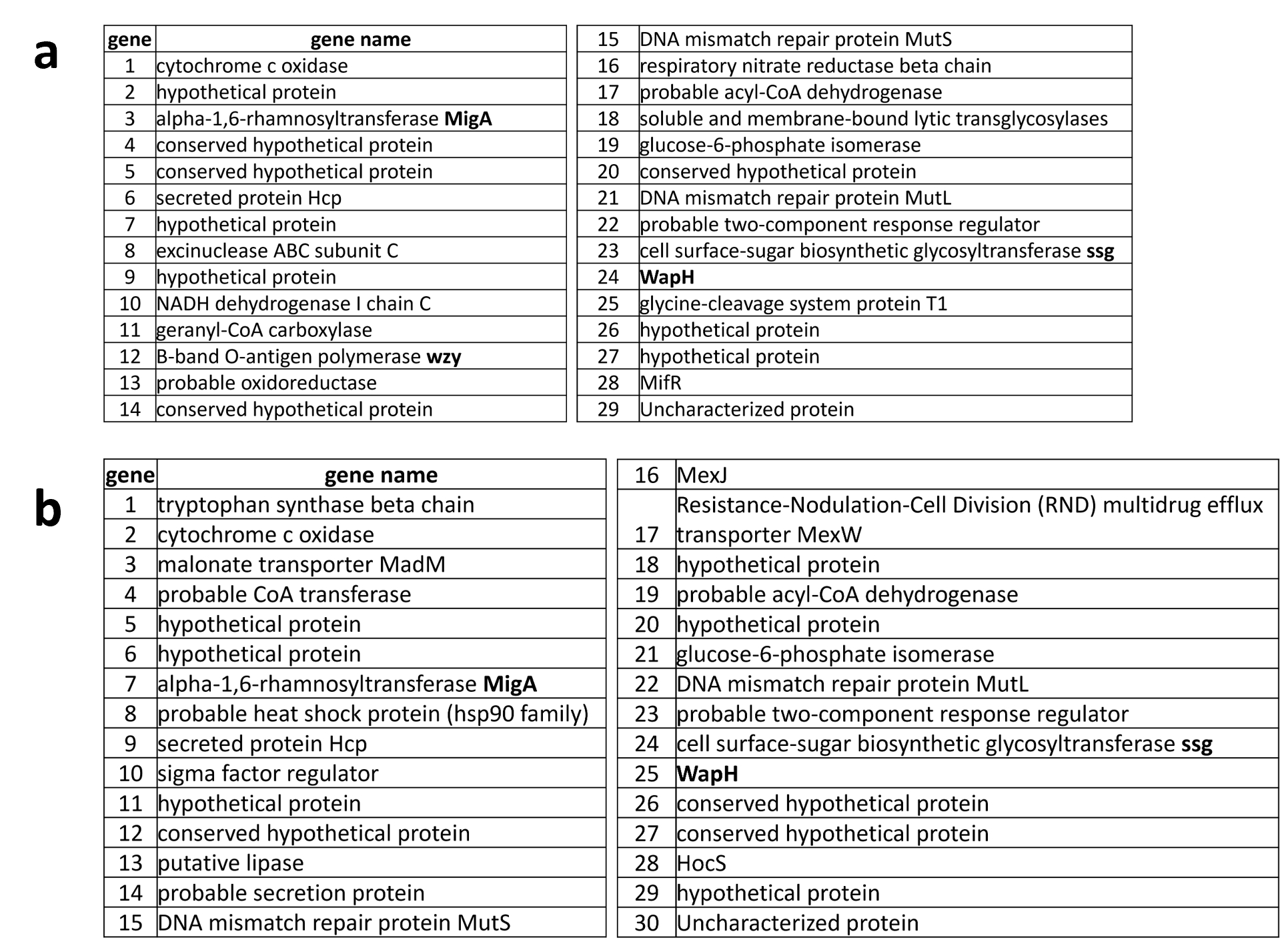

**Supplementary Figure 4**: **Identification of Genes associated with LPS-5 Phage Infection through Transposon Mutagenesis Experiments.** Full legends to the volcano plots in Figure 5A and Supplementary Figure 5A for LPS-5 at MOI 0.1 (a.) and MOI 20 (b.). Volcano plot describes genes with a significant difference in counts (fold change) between conditions with and without LPS-5 phage.

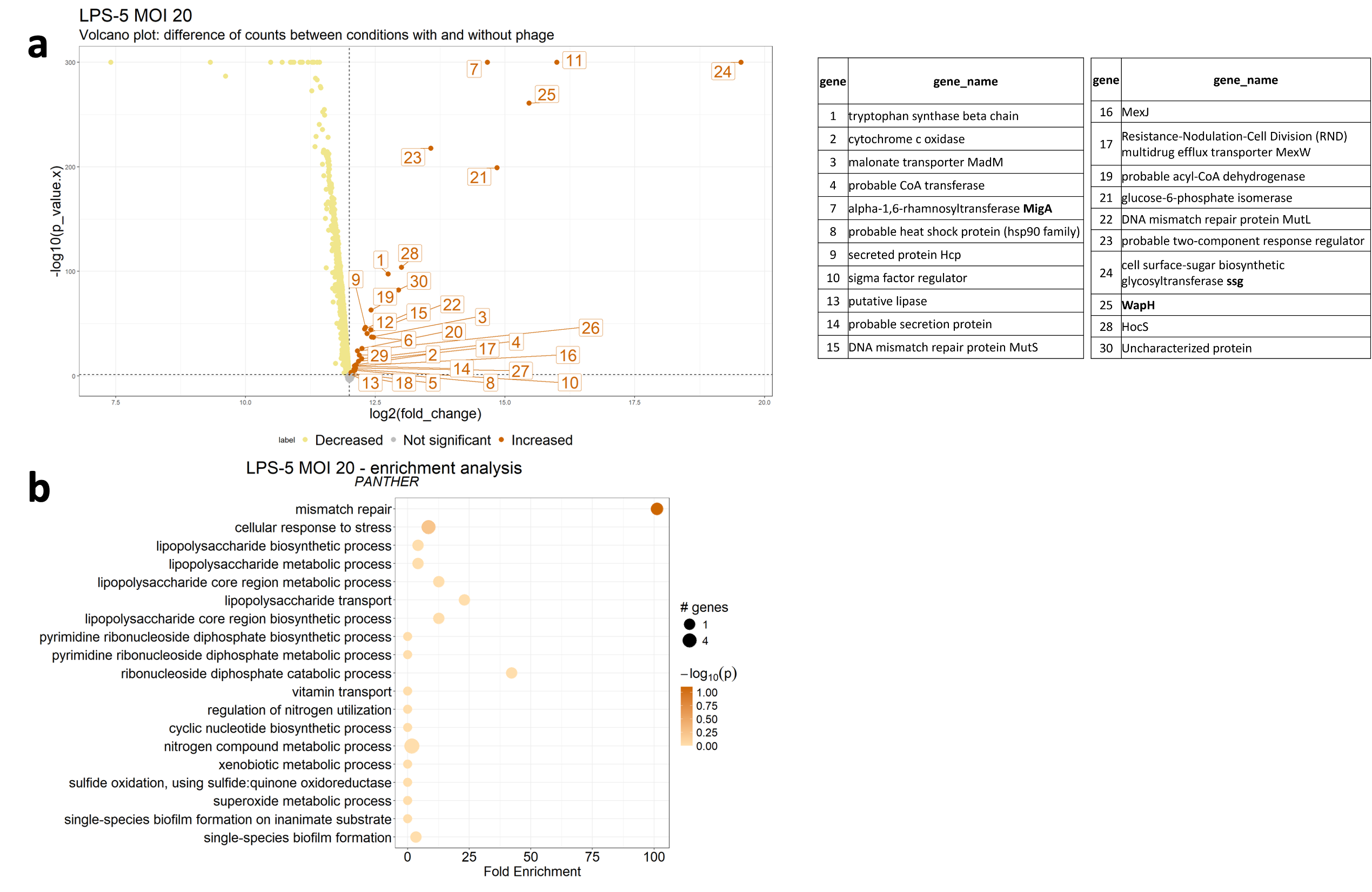

**Supplementary Figure 5**: **Identification of Genes associated with LPS-5 Phage Infection through Transposon Mutagenesis Experiments a,** This volcano plot describes genes with a significant difference in counts (fold change) between conditions with and without LPS-5 phage at MOI 20. Like the previous plot, orange signifies a significant increase in gene counts, yellow stands for a significant decrease, and gray represents non-significant genes. **b,** Similar to the previous functional analysis plot, this section summarizes the functional enrichment analysis results for the LPS-5 phage at MOI 20 with PANTHER.

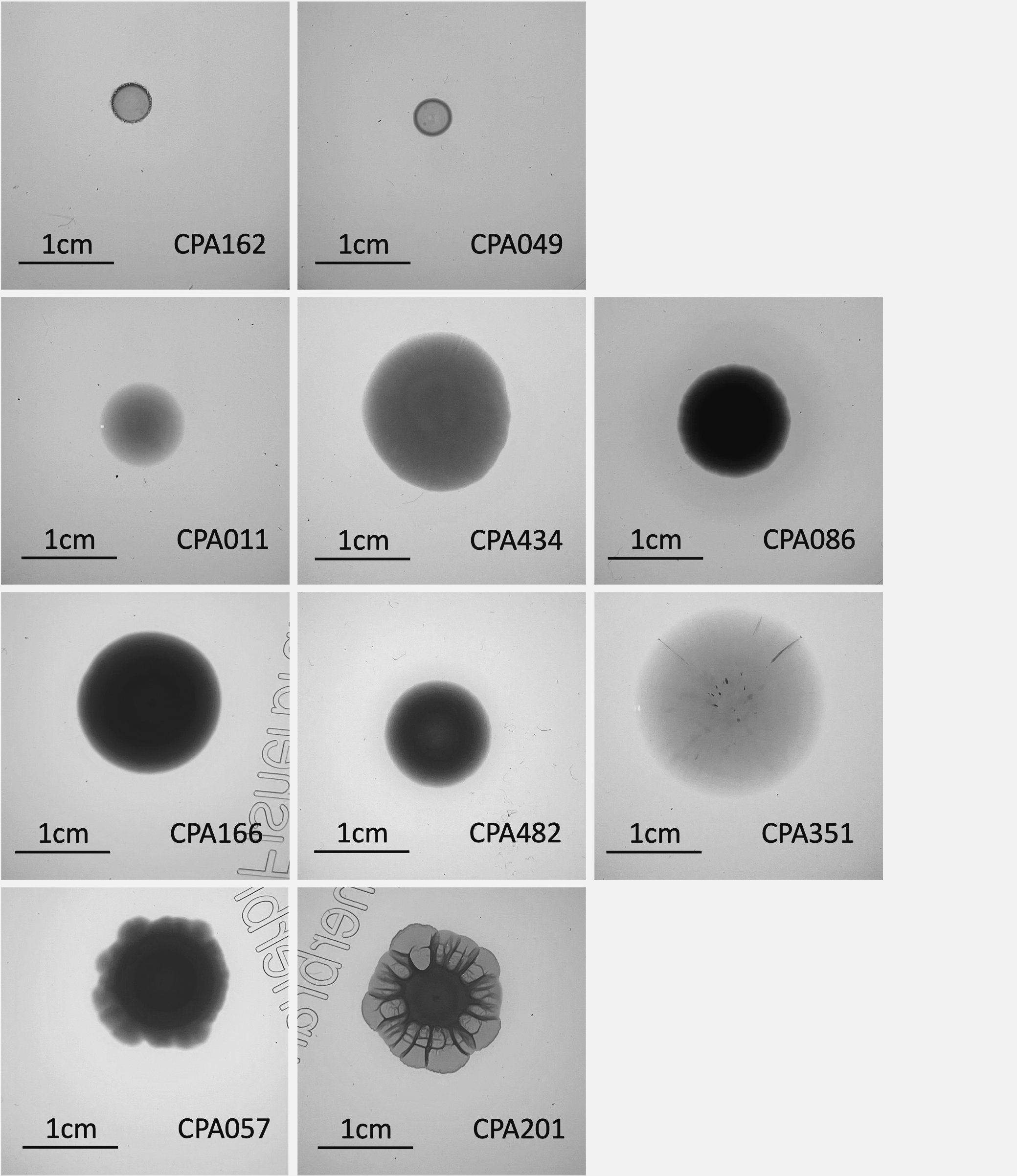

**Supplementary Figure 6: O-AG/LPS morphology of the clinical isolates.** The function of the O-antigen in clinical isolates was assessed using a rugae O-AG/LPS assay. Several bacterial strains did not form a colony biofilm rugae (rough colony morphotype), suggesting mutations in their O-antigen within the LPS.

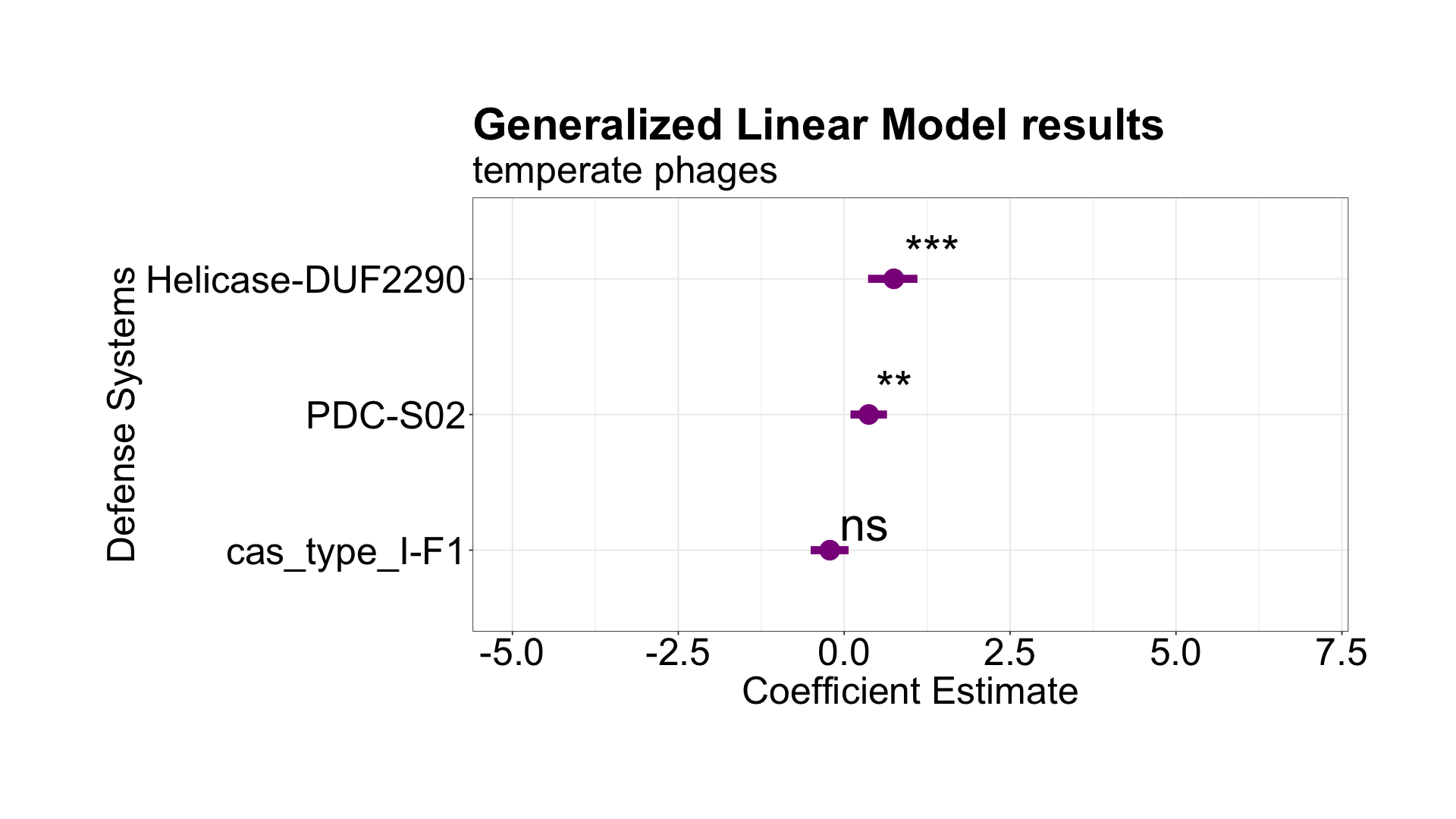

**Supplementary Figure 7: Bacterial Defense Systems Correlate with Temperate Phage Prevalence.** Bacterial defense systems are associated with the number of temperate phages in the bacterial genome, even after removal of systems encoded within temperate phage islands in the bacterial genome.

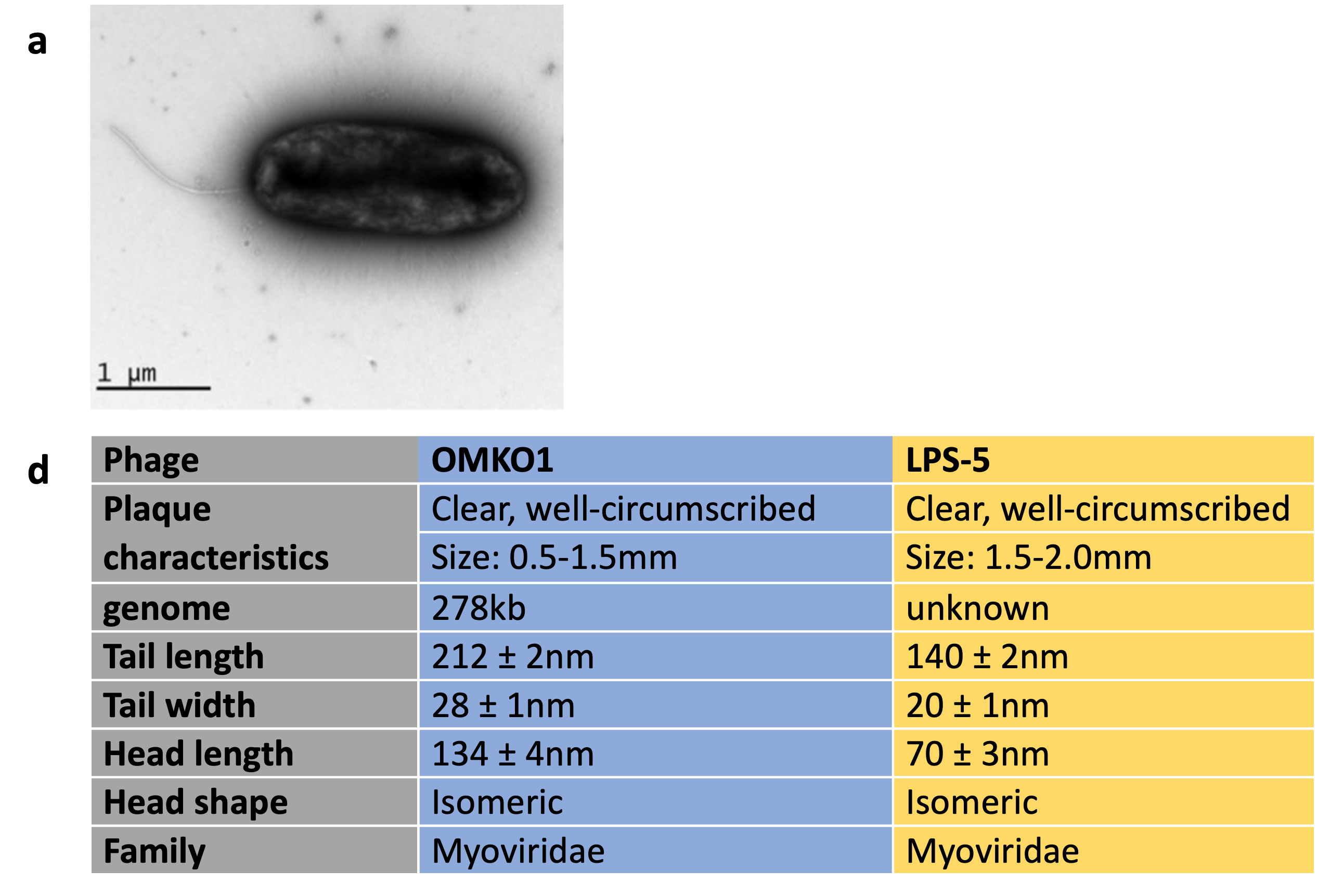

**Supplementary Figure 8**: **Bacteriophages OMKO1, LPS-5 and their host Pseudomonas aeruginosa.** **A,** TEM image of *Pseudomonas aeruginosa*, the natural host of phages OMKO1 and LPS-5. **B,** Characteristics of phages OMKO1 and LPS-5.
